# Genetic Variation, Iron Status, and FGF23 Signaling Converge to Regulate Renal Calcium Buffering in Sickle Cell Disease

**DOI:** 10.64898/2026.09.23.752598

**Authors:** Grace Xie, Kathryn Janeczko, Ella Williams, Yueh-Ting Wang, Rafiou Agoro

## Abstract

Sickle cell disease (SCD) causes heterogeneous mineral imbalances including variable degrees of hypocalcemia. The kidney controls systemic calcium by reabsorbing calcium from the glomerular filtrate via paracellular transport and transcellular transport in the nephron tubules, yet it is unknown whether these processes are modulated by genetic or environmental factors or disrupted in SCD. Using SCD mouse models and single-cell multiomics, we identify the distal convoluted tubule (DCT) as the nephron segment most susceptible to calcium reabsorption dysfunction in SCD, mainly via reduction of calcium buffer protein calbindin 1 (CALB1). We show that CALB1 and its encoding mRNA are decreased in DCT cells in SCD, alongside decreased Klotho (KL)-dependent fibroblast growth factor (FGF) 23 signaling and intracellular calcium signaling. Dietary iron restriction reduces CALB1, KL, and calcium exporter SLC8A1 levels in SCD kidneys. Loss of CALB1 shifts DCT cells toward energy-inefficient glycolysis with the metabolite 2,3-diphosphoglycerate impairing KL-dependent FGF23 signaling to create a feed-forward loop suppressing calcium reabsorption. Analysis of gene expression and protein quantitative trait loci data from kidneys of genetically diverse mice revealed that *Calb1* expression levels are highly heritable and co-regulated with *Slc8a1*, identifying a genetic axis that dictates differential capacities for calcium buffering and blood export in the kidney. Together, these findings support a model in which genetic variation, dietary iron status, and FGF23 signaling converge on DCT calcium buffering to reduce renal calcium reabsorption in the SCD kidney. This points to personalized, genotype- and iron-dependent strategies for managing mineral metabolism in SCD patients.

## Introduction

Sickle cell disease (SCD) is a monogenic Mendelian disease with clinical feature heterogeneity arising from a combination of variable genetic and environmental factors.^1,2^ Some patients are repeatedly hospitalized with SCD symptoms, whereas others have few symptoms and less vaso-occlusive events.^3–5^ Hypocalcemia is frequent in SCD, worsens during acute vaso-occlusive events, and elevates mortality risk,^6^ although some patients do not develop hypocalcemia at all.^7^ How genes and the environment interact to control calcium homeostasis in SCD remains unclear. In healthy individuals, blood calcium levels are affected by interrelated biological mechanisms across multiple organs, including intestinal calcium absorption, calcium release from bone resorption, and calcium reabsorption from kidney glomerular filtrate.^8^ In the kidney, calcium homeostasis is controlled at different nephron sites. In the proximal tubule (PT) and loop of Henle, calcium is passively reabsorbed paracellularly through claudin-2 and -16 tight junctions.^9,10^ In the distal tubule, which comprises the distal convoluted tubule (DCT) and connecting tubule (CNT), calcium is actively reabsorbed transcellularly through transient receptor potential vanilloid (TRPV) 5.^10,11^ Active calcium reabsorption in the distal tubule is positively regulated by fibroblast growth factor 23 (FGF23), parathyroid hormone (PTH), and calcitriol (1,25-dihydroxyvitamin D; VitD).^12^ The levels and activity of these circulating molecules are shaped by both genetic variation and environmental factors such as diet, geographic location, and multiple disease states.^8,13,14^ Understanding how genetic modifiers of these regulatory pathways interact with environmental factors to alter renal calcium handling may therefore explain the heterogeneous development of hypocalcemia observed in SCD patients.

Here, using mouse models, we reveal that the DCT is the nephron segment where calcium-handling pathways are most reprogrammed in SCD. SCD decreased the abundance of calcium-buffering gene transcripts and proteins in the DCT, with the calcium-binding protein calbindin 1 (CALB1) being co-downregulated alongside the FGF23 co-receptor protein, Klotho (KL). Dietary iron restriction further decreased CALB1 and KL protein levels in the kidney, indicating that iron deficiency compounds the effects of SCD. Mechanistically, the SCD-associated decrease in calcium buffering capacity drove DCT cells toward an energy-inefficient glycolytic state, leading to accumulation of 2,3-diphosphoglycerate (2,3-DPG), a metabolite that we demonstrate herein reduces KL-dependent FGF23 signaling, a canonical driver of the kidney’s transcellular calcium reabsorption pathway.^15^ Using kidney data from genetically diverse mice, we found specific gene alleles linked to high heritability of calcium-buffering capacity, underscoring a strong contribution of genetic variation to kidney calcium handling. Together, these findings show that heritable factors, dietary iron status, and hormonally mediated, KL-dependent FGF23 signaling converge to control calcium handling in the DCT, shaping both DCT cell metabolism and the molecular pathways governing calcium reabsorption.

## Results

### SCD impairs kidney calcium buffering capacity in distal tubule cells

To determine the degree of heterogeneity in circulating calcium levels across SCD patients, we conducted a meta-analysis of existing clinical data from SCD patients in different countries published before July, 2026. Of six case-versus-control studies (from 1987 to 2005) conducted in Nigeria (one study in Lagos and one in Port Harcourt), Sudan, Ghana, Iraq, and Turkey,^16–21^, five reported lower serum calcium levels in SCD relative to controls, with only the Bayazit *et al*.^21^ study in Turkey reporting non-significant higher levels. A random-effects (DerSimonian–Laird) meta-analysis pooling yielded a significant negative overall effect of SCD on calcium levels (Hedges’ g = −0.71) with a 95% CI ranging from −1.26 to −0.17 (**Figure 1a**). These findings provide evidence that SCD is associated with low average serum calcium and that inter-geographical variation confounds our ability to understand the relationship between calcium homeostasis and SCD. This underscores the need to consider how genetic and environmental factors drive calcium level variability in the context of SCD.

**Figure 1:**
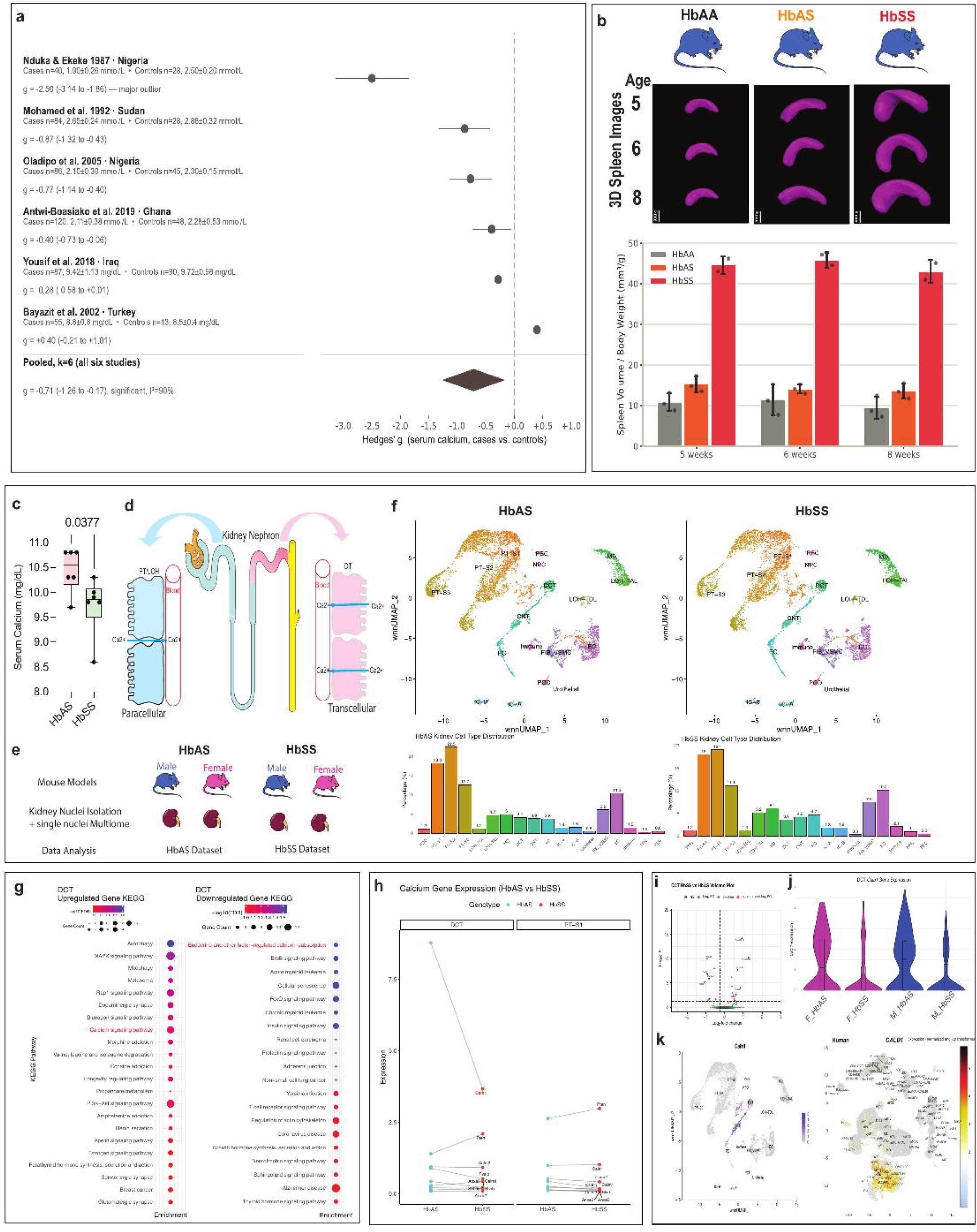
Transcriptional reprogramming of Calb1 in SCD. (**a**) Forest plot of random-effects meta-analysis of six studies reporting serum calcium in SCD patients vs. healthy controls. Circles and lines show individual-study Hedges’ g with 95% confidence intervals, respectively; the diamond shows the pooled random-effects (DerSimonian–Laird) estimate across all six studies (g = −0.71, 95% CI −1.26 to −0.17, I² = 90%). Studies are ordered by effect size, from most negative (lower calcium in SCD cases) to most positive. Country of origin is indicated for each study. Negative values indicate lower serum calcium in SCD cases relative to controls; the dashed vertical line marks no difference (g = 0). (**b**) HbAA, HbAS, and HbSS mice were MicroCT scanned at ages 5, 6, and 8 weeks to generate 3D spleen images from which body-weight normalized spleen volumes were calculated. (**c**) Decreased serum calcium in male HbSS vs. HbAS. (**d**) Diagram of renal pathways for calcium reabsorption into the bloodstream. (**e**) Overview of multiomics analysis. Single cells from HbAS and HbSS pooled male and female kidneys were subjected to scRNA-seq and downstream computational analysis was performed to deconvolute male and female cells. (**f**) Unsupervised uniform manifold approximation and projection (UMAP) clustering identified 18 distinct renal cell types (top). Summary bar plots of kidney cell type distribution (bottom). (**g**) Dot plot of KEGG pathway enrichment in DCT of HbSS vs. HbAS. (**h**) Pair plots comparing expression levels of calcium handling-related genes between HbAS and HbSS. (**i**) Volcano plots of differentially upregulated and downregulated genes in DCT highlight significant downregulation of Calb1. (**j**) Violin plots show Calb1expression in the DCT in HbSS vs. HbAS. (**k**, left) Feature plot of Calb1 expression in control HbAS mouse kidney pinpoints its restricted expression to distal tubule segments of the nephron (DCT: distal convoluted tubule; CNT: connecting tubule). (**k**, right) Feature plot of healthy human kidney scRNA-seq data^24^ highlighting CALB1 expression in DCT cells.

Because impairment of renal calcium reabsorption reduces circulating calcium,^22^ we sought to determine if this homeostatic reabsorption process is altered in SCD kidneys. For this, we used the humanized Townes mouse model^23^ in which mice produce only human hemoglobins (Hb), including normal healthy beta-globin (HbA) or SCD beta-sickle-globin (HbS). This model mimics the human pathophysiology of SCD (HbSS mice), sickle cell trait (HbAS mice), and healthy phenotypes (HbAA mice). Like human SCD patients, HbSS mice developed hemolytic anemia linked with extramedullary erythropoiesis in the spleen and spleen enlargement,^23^ a classical marker of SCD. In comparison to either HbAA or HbAS, HbSS mice exhibited spleen enlargement at 5, 6, and 8 weeks (**Figure 1b**), while spleen volume was not different between HbAS and HbAA (**Figure 1b**). Biochemical profiling of male HbSS mice revealed low blood calcium compared to HbAS (**Figure 1c**), recapitulating the hypocalcemic shift observed in patients and validating the Townes model as a suitable system for investigating its mechanistic basis.

To identify potential SCD-associated changes in renal calcium reabsorption pathways across nephron cell types, we performed single-cell multiomics of kidneys from male and female HbSS vs. HbAS mice (**Figure 1e**). To both identify nephron segment-specific cell types and uncover changes in their expression and regulation of calcium handling-related genes, we performed single-nuclei RNA sequencing (snRNA-seq) and single-nuclei ‘assay for transposase-accessible chromatin’ sequencing (snATAC-seq) (data quality control in **Supplementary Figure 1a**). Unsupervised uniform manifold approximation and projection (UMAP) clustering of the RNA-seq data identified 18 kidney cell types in male and female mice, including epithelial, endothelial, and immune cells (**Figure 1f** and **Supplementary Figure 1b**). As expected, epithelial PT cells were the most represented population (50.93%) (**Figure 1f**). Differential gene expression and KEGG pathway analysis of epithelial PT-S1 and DCT cells revealed that, in the DCT, the “Calcium signaling” pathway was upregulated in HbSS relative to HbAS in both male and female mice, while the “Endocrine-regulated calcium reabsorption” pathway was downregulated (**Figure 1g**), predicting an increase in intracellular calcium alongside disruption of hormone-dependent calcium import/export mechanisms. No changes in calcium reabsorption pathways were observed in PT-S1 (**Supplementary Figure 1c**). Comparing transcript abundance of calcium handling-related genes in HbSS vs. HbAS mice revealed a marked decrease in *Calb1* specifically in the DCT, the nephron segment responsible for hormone-dependent, transcellular calcium reabsorption (**Figure 1h**). This was observed in both male and female mice (**Figure 1j**). *Calb1* encodes CALB1, a protein that buffers intracellular calcium levels by binding calcium ions and facilitates their trafficking to the plasma membrane for export. Other calcium buffer-encoding genes, including those for calbindin 3 (S100G) and parvalbumin (PVALB), showed modest decreases in HbSS vs. HbAS (**Supplementary Figure 1D**). In both healthy mouse and human kidney^24^, *Calb1* and *Pvalb* were found to be highly expressed in DCT and CNT cells (**Figure 1k and Supplementary Figure 1e**). Together, these findings indicate that SCD disrupts calcium homeostasis specifically by impairing calcium buffering capacity in distal tubule cells.

### FGF23 controls renal expression of the calcium buffer gene Calb1

To understand how expression of *Calb1* levels might be altered in the SCD kidney, we first searched for cell type-specific changes in chromatin accessibility associated with *Calb1* expression level. In the kidney of HbAS control mice, *Calb1* expression in DCT and CNT cells was associated with a chromatin accessibility peak (chr4:15,880,814–15,881,806) in the promoter region (**Figure 2a**). *Calb1* expression was downregulated in DCT cells in HbSS, concomitant with decreased signal at the chromatin accessibility peak. To determine whether differentially accessible chromatin in HbSS vs. HbAS DCT cells is associated with known *cis*-regulatory elements at the *Calb1* promoter, we scanned the chr4:15,880,814–15,881,806 sequence with JASPAR.^25^ This identified a high-confidence repressive element 1 (RE1) binding-site motif for the hypoxia-responsive^26^ RE1-silencing transcription factor (REST) at chr4:15,881,291–15,881,311 (+ strand; 5′-AGCAGCACCGCGGACAGCGCC-3′), scoring 92.4% of the maximum possible position weight matrix (PWM) score and independently confirmed by the UCSC JASPAR track (score = 883) (**Figure 2b**). This site lies within the 5′ untranslated region (UTR) between the annotated transcription start site (TSS at 15,881,263) and the translation start site (cdsStart at 15,881,398) (**Figure 2c**). Cross-species comparison to publicly available human kidney single-nucleus (sn) ATAC-seq data for 237,274 single cells from 30 male and female kidneys^27^ revealed that the promoter region of human *CALB1* likewise contains a chromatin accessibility peak harboring a predicted RE1 element in DCT cells (**Supplementary Figure 2a**), suggesting that REST-mediated transcriptional repression of *CALB1* is conserved between mouse and human. Further, there was enrichment of the REST motif in snATAC-seq data of DCT cells from HbSS vs. HbAS mice (**Supplementary Figure 2b**).

**Figure 2:**
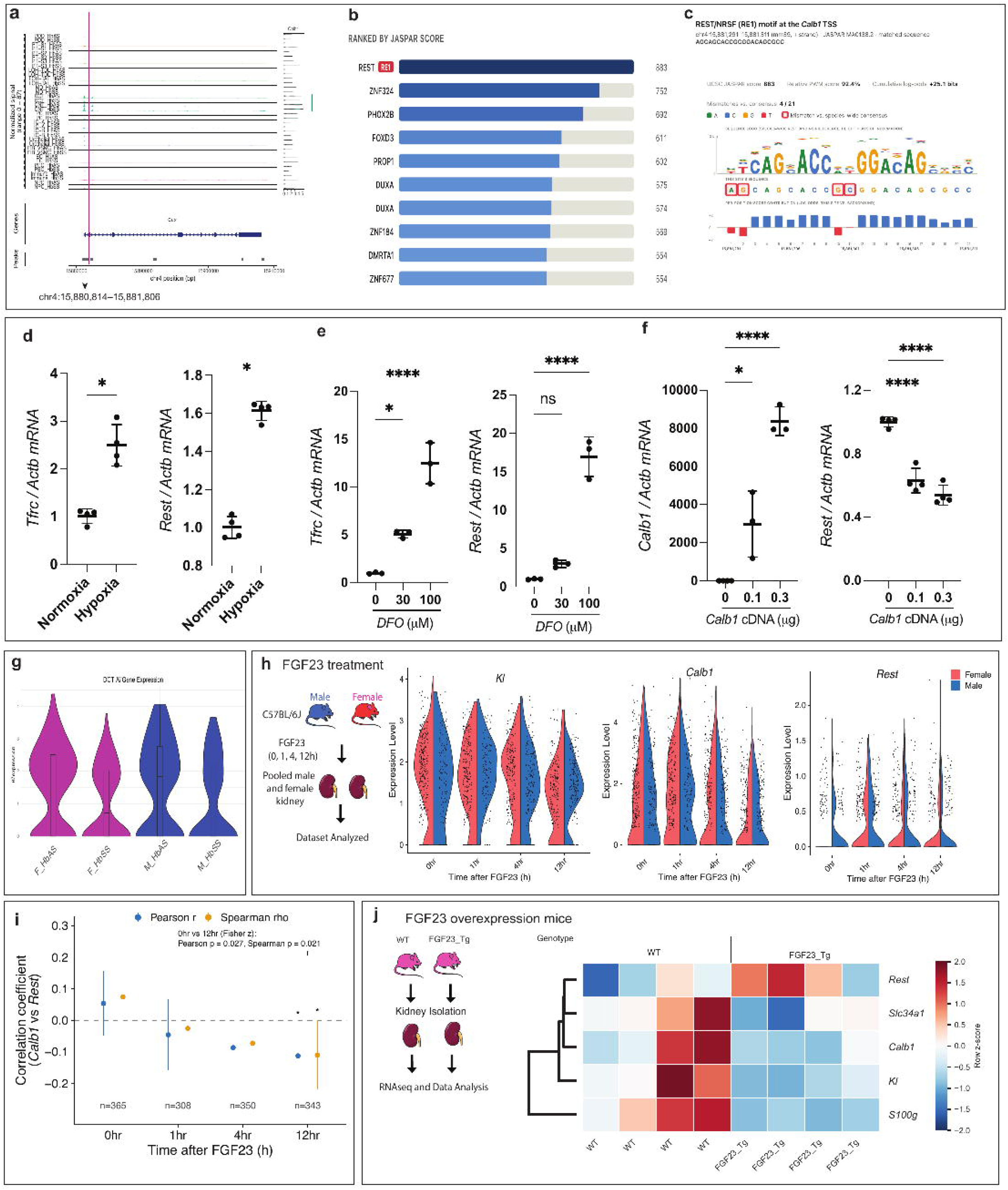
Calb1 is regulated by the hormone FGF23 in DCT cells. (**a**) Left plot: Peaks of Calb1 locus chromatin accessibility peaks (green) from snATAC-seq analysis, spanning chr4:15,880,814–15,881,806 which is located within the magenta box, close to the promoter region (GRCm38/mm10). Right: RNA levels in HbSS vs. HbAS pooled male and female kidneys from snRNA-seq analysis shown as horizontal violin plot. (**b**) Ranking of transcription factor binding-site motifs within the Calb1 chromatin accessibility region using the JASPAR database. (**c**) The REST sequence logo derived from the JASPAR position frequency matrix, highlighting that the site scores in the top 8% of the matrix (92.4% of the maximum possible PWM score). (**d**) DCT cell line 209 was plated for 32 h followed by 16 h incubation in hypoxic (1% O_2_) or normoxic conditions (21% O_2_) and Tfrc, and Rest mRNAs quantitated by RT-qPCR. (**e**) DCT cell line 209 treated with deferoxamine at different concentrations or vehicle alone as a control for 16 h followed by RT-qPCR analysis of Tfrc and Rest. (**f**) DCT cell line 209 transfected with Calb1 cDNA-expressing plasmid (0.1 μg and 0.3 μg) or empty expression plasmid (0.3 μg pCMV6) as a control for 32 h followed by 16 h in hypoxic (1% O_2_) conditions and Calb1, Tfrc, and Rest mRNAs quantitated by RT-qPCR. (**g**) Violin plots show Kl expression in the DCT of HbAS vs. HbS male and female mice. (**h**) Single-cell RNA-seq analysis of kidney tissue from C57BL/6J mice treated with FGF23 (0 h, 1 h, 4 h, and 12 h). Violin plots illustrate the expression levels of *Kl, Calb1*, and *Rest* within the distal convoluted tubule (DCT) cell population. Blue and red colors denote male and female datasets, respectively. (**i**) Calb1–Rest correlation across the FGF23 time course in DCT cells. Pearson (blue) and Spearman (orange) correlation coefficients between Calb1 and Rest expression in DCT cells at 0, 1, 4, and 12 h post-FGF23 (n = 365, 308, 350, 343 DCT cells). Error bars: 95% CI (Fisher z). *p < 0.05. Bracket: direct 0 h vs. 12 h comparison (Fisher z-test for independent correlations; Pearson p = 0.027, Spearman p = 0.021). (**j**) Kidneys from 8-week-old female mice expressing a FGF23 transgene were subjected to RNA-seq. Heatmap of row-scaled (z-score) normalized expression for Kl, Slc34a1, Calb1, S100g, and Rest in WT (n=4) and FGF23_Tg (n=4), hierarchically clustered by gene (average linkage, Euclidean distance). Kl, Slc34a1, Calb1, and S100g were reduced in FGF23_Tg; Rest shows the opposite trend.

To test whether hypoxia directly regulates *Rest* expression in kidney DCT cells, we placed the DCT cell line 209 in normoxic and hypoxic conditions and measured the expression of *Rest* and *Tfrc*, a canonical hypoxic marker that encodes transferrin receptor protein 1 (TFRC). Hypoxia induced *Tfrc* and *Rest* in 209 cells (**Figure 2d**). Since iron deficiency impairs oxygen transport^28^ and promotes hypoxia (and HIF stabilization), we treated 209 cells with different concentrations of the iron-chelating drug deferoxamine (DFO) to orthogonally test whether restricting iron availability regulates *Tfrc* and *Rest.* The resulting iron deficiency increased expression of both genes, and importantly, the effect was dose-dependent (**Figure 2e**). Further, overexpression of *Calb1* suppressed *Rest* expression (**Figure 2f**), suggesting an inverse relationship between *Calb1* and *Rest* expression in DCT cells.

Next, we hypothesized that *Calb1* expression is influenced by alteration in signaling pathways associated with the calcium-regulating hormones PTH and VitD or the bone-derived mineral homeostasis-regulating hormone FGF23, as genes encoding the receptors of these hormones (*Pth1r*, *Vdr*, *Fgfr1*, and *Kl*) are expressed in mouse and human DCT cells (**Supplementary Figure 2c**). Among these receptors, only *Kl* was significantly downregulated in DCT cells in HbSS vs. HbAS (**Figure 2g**), whereas *Pth1r*, *Vdr*, and *Fgfr1* did not show significant changes (**Supplementary Figure 2d**). We did not find enrichment of VitD receptor (VDR) binding site motifs in snATAC-seq data from HbSS vs. HbAS DCT cells (**Supplementary Figure 2e**). Further, circulating PTH was not elevated; rather, PTH levels were decreased in HbSS vs. HbAS male and female mice (**Supplementary Figure 2f**). These findings suggest that VitD may not directly contribute to transcriptional reprogramming of *Calb1* in the DCT of SCD kidneys.

Because we did not observe an enrichment of VDR binding-site motifs at the *Calb1* promoter region and the mechanism of *Calb1* regulation is likely PTH-independent in DCT cells as there was no increase in circulating PTH, we hypothesized that *Calb1* is regulated by another calcium-regulating hormone, FGF23. To test this hypothesis, we analyzed our previously published kidney single-cell data^29^ from a time course of male and female C57BL/6J mice following FGF23 treatment (**Figure 2h**). *Calb1* was downregulated in DCT cells at 4 and 12 h post-treatment whereas *Rest* was upregulated (**Figure 2h**), suggesting a potential inverse regulatory effect that is influenced by FGF23 (**Figure 2j**). Further, kidneys from a mouse model of chronically elevated FGF23 levels (FGF23-TG mice)^30^ showed decreased expression of *Slc34a1*, a classical marker of FGF23 activity, as well as of *Kl*, *Calb1*, and *S100g*, contrasting the increased expression of *Rest* (**Figure 2j**). Taken together, these data are consistent with a model in which FGF23 and hypoxia increase REST levels to downregulate *Calb1* expression in the DCT.

### KL-dependent FGF23 signaling controls renal calcium buffering targets in kidney dysfunction

In the SCD state (7 weeks of age), bone and bone marrow expression of *Fgf23* is upregulated in female HbSS vs. HbAA mice (**Figure 3a**). This change is associated with increased levels of circulating C-terminal FGF23 peptide (C-term FGF23; **Figure 3a**). In parallel, immunoassays revealed a decrease of KL in kidneys of HbSS vs. HbAS mice (**Figure 3b-c**). To determine if high FGF23 and low KL levels coordinately regulate *Calb1,* we leveraged *in vivo* mouse models of chronic kidney disease (CKD) and acute kidney injury, which both exhibit elevated FGF23.^31^ To induce CKD in mice, we fed male and female C57BL/6J mice an adenine diet (AD) for 2 and 4 weeks (**Figure 3d**). Compared to mice on the casein control diet (CAS), there was a progressive increase in blood urea nitrogen (BUN) levels in AD-fed mice, indicating a CKD state (**Supplementary** Figure **3a**). Levels of circulating intact and C-terminal FGF23 increased after AD treatment (**Figure 3e**). We then hypothesized that *Calb1* is downregulated in kidneys of CKD-state mice concomitant with reduced levels of *Kl* in CKD.^32,33^ To test this, we performed kidney bulk RNA-seq of male and female mice fed AD vs. CAS. Principal component analysis (PCA) reflected induction of the CKD state in AD-fed mice when compared to control CAS-fed mice (**Supplementary** Figure **3b**). We found decreased renal expression of *Calb1* and *Kl* in mice fed AD vs. CAS (**Figure 3f**). In addition to *Calb1,* other calcium buffer-encoding genes, *S100g* and *Pvalb*, were downregulated (**Figure 3f**).

**Figure 3:**
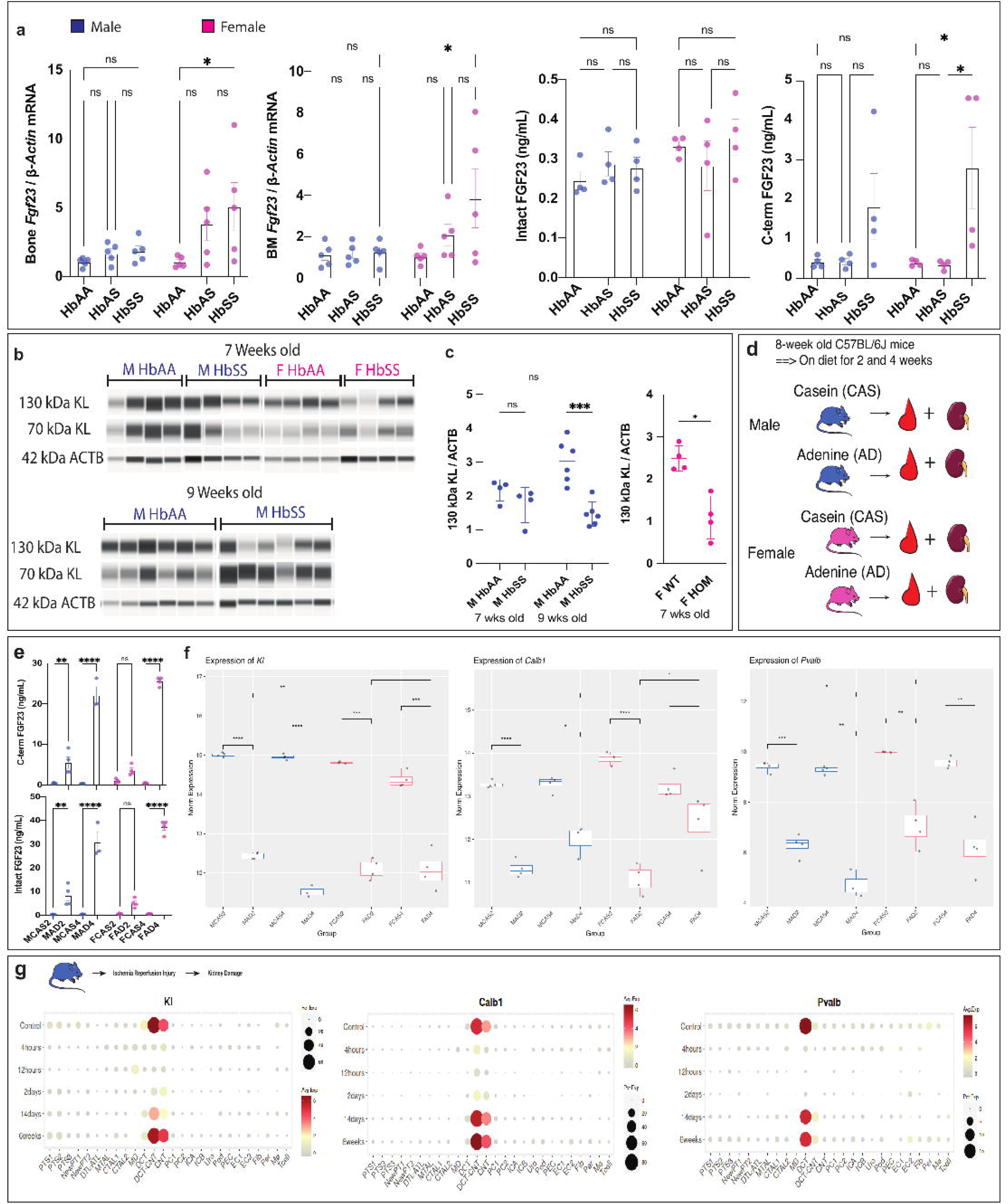
KL-dependent FGF23 signaling controls Calb1 in acute and chronic kidney dysfunction. (**a**) Bone and bone marrow from male and female HbAA, HbAS, and HbSS mice were subjected to RT-qPCR analysis of Fgf23 gene expression (top) and ELISA quantification of circulating intact or C-term FGF23 (bottom). (**b-c)** Immunoassay of KL in kidney (**b**) with levels of each isoform quantified relative to β-actin (**c**). (**d**) Overview of CKD mouse study. (**e**) Circulating intact and C-term FGF23 levels were measured by ELISA in 8-week-old male and female C57BL/6J mice fed a casein control diet (CAS) or CKD-inducing adenine diet (AD) for 2 and 4 weeks. (**f**) mRNA levels for Kl, Calb1, and Pvalb in kidneys of male and female C57BL/6J mice after 2 or 4 weeks on CAS or AD. (**g**) Kl, Calb1, and Pvalb expression during acute kidney injury in 8–10-week-old male C57BL/6J mice.

To test the concept that KL-dependent *Calb1* regulation occurs in a specific segment of the distal tubule, we used a scRNA-seq dataset from mice that underwent bilateral ischemia–reperfusion injury (IRI),^34^ a model of acute kidney injury. The male mouse kidneys were analyzed at 4 h, 12 h, 2 days, 14 days, and 42 days after IRI (see UMAP in **Supplementary Figure 3c**) and exhibited strong induction of the kidney injury marker *Kim1* (also known as *Havcr1*) in PT cells at 12 h that fell by 2 days (**Supplementary Figure 3d**). The onset of injury produced a marked decrease in *Kl, Calb1*, and *Pvalb* expression in the DCT and CNT at 4 h before recovering at 42 days (**Figure 3h**). Taken together, our findings suggest a scenario in which kidney dysfunction causes cells of the distal tubule to become unresponsive to circulating FGF23 due to the drop in distal tubule *Kl* expression, which is then accompanied by a decrease in *Calb1* and *Pvalb* expression. We speculate that this scenario is relevant to kidney function in the context of calcium reabsorption impairment in SCD and that key environmental factor may regulate this pathway.

### Iron restriction promotes kidney hypoxia and decreases KL and calcium buffering proteins in SCD

Because renal hypoxia is a common occurrence in CKD and acute kidney injury,^35^ we hypothesized that environmental factors that promote renal hypoxia would also reduce the levels of calcium buffers and exacerbate renal calcium buffering impairment in SCD. Since iron deficiency impairs oxygen transport,^28^ we reasoned that an iron restriction regimen would mimic hypoxia. Thus, we fed male HbAA and HbSS mice iron-replete control (IronCtrl) and iron-deficient (IronDef) diets for four weeks (**Figure 4a**). As expected, circulating ferritin levels decreased in HbAA_IronDef vs. HbAA_IronCtrl mice (**Figure 4b**), suggesting tissue iron depletion. As expected, circulating ferritin was higher in HbSS_IronCtrl vs. HbAA_IronCtrl mice (**Figure 4b**). This increased level is likely because of an elevated inflammatory state and/or iron sequestration in tissues (**Figure 4b**). Circulating total FGF23 (C-term FGF23) levels were also higher in HbSS_IronCtrl vs. HbAA_IronCtrl mice (**Figure 4c**).

**Figure 4:**
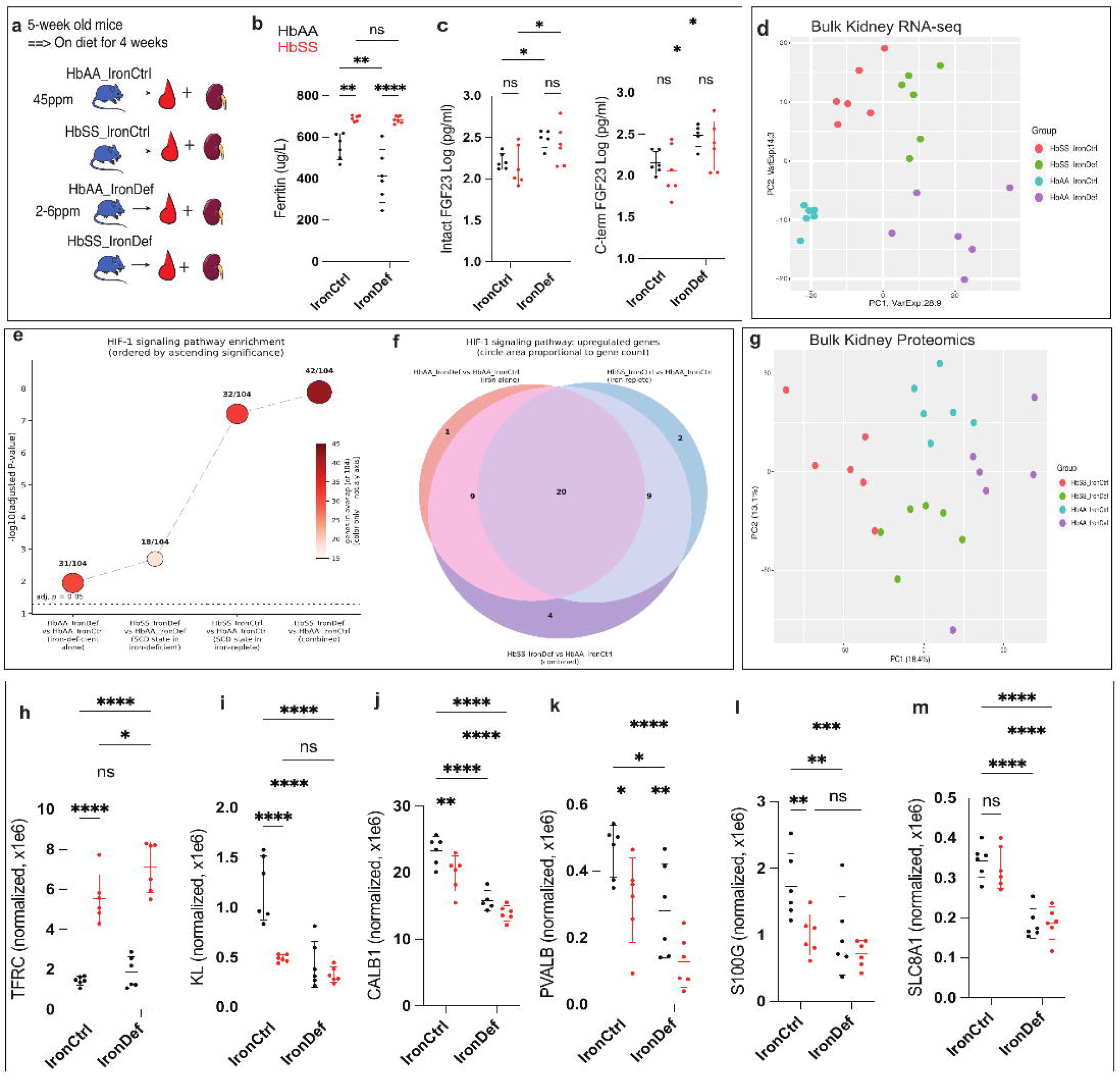
Iron-mediated transcriptional reprogramming of Calb1 in the mouse kidney. (**a**) Experimental overview. Male HbAA and HbSS mice were fed iron-replete (IronCtrl) or iron-deficient (IronDef) diets for four weeks followed by analyses of blood plasma and kidneys. (**b**) Ferritin levels in plasma were measured using a colorimetric assay. (**c**) Circulating levels of intact and C-term FGF23 quantified by ELISA of plasma from HbAA and HbSS mice on the IronCtrl or IronDef diets. (**d**) PCA plot of male HbAA and HbSS kidney bulk RNA-seq data. (**e**) Pathway enrichment analysis of that exhibited expression-level changes in the SCD or dietary states identified 104 genes known as upstream or downstream regulators of the HIF-1α signaling pathway. (**f**) Bubble plot of these HIF-1α signaling pathway-associated genes highlights that most of them are driven by both the SCD and dietary states, with a subset exhibiting an additive effect on expression. (**g**) PCA plot of bulk kidney proteomics data from male HbAA and HbSS kidney. (**h-m**) TFRC, KL, CALB1, PVALB, S100G, and SLC8A1 protein abundances measured by mass spectrometric analysis.

To find molecular pathways in the kidney that are reprogrammed by the SCD state in the context of dietary iron restriction, we performed bulk RNA-seq. PCA of the data revealed distinct clustering of HbSS samples and IronDef samples (**Figure 4d**). Pathway enrichment analysis revealed an enrichment of genes associated with the HIF-1α signaling pathway in the SCD state of HbSS_IronCtrl mice vs. HbAA_IronCtrl mice (**Figure 4e**). Changes for a subset of these genes were further pronounced under iron restriction in HbSS_IronDef mice vs. HbAA_IronCtrl mice (**Figure 4e-f**). To determine the impact of iron restriction on calcium binding potential in the kidney, we performed kidney bulk proteomics. PCA of the proteomics data revealed distinct clustering of HbSS samples and IronDef samples (**Figure 4g**). In HbSS, there was an increase in abundance of the hypoxia marker TFRC, and this increase was promoted by dietary iron restriction (**Figure 4h**). This additive effect of SCD and iron restriction on TFRC suggests an additive effect on HIF-1α signaling activity when iron restriction is coupled with SCD.

In the SCD state and the dietary iron restriction state, kidney KL was decreased (**Figure 4i**). Further, we found that iron restriction decreased the abundance of calcium buffers, including CALB1, PVALB, and S100G, in the kidney of HbAA mice (**Figure 4j-l**). Of note, restricting iron in HbSS mice further decreased the abundance of CALB1, S100G, and PVALB when compared to either iron restriction or the SCD state alone (**Figure 4j-l**). Finally, iron restriction decreased calcium exporter SLC8A1 and ATP2B4 levels (**Figure 4m** and **Supplementary Figure 4a**). Thus, a low-iron status that promotes hypoxia and decreases renal KL levels in SCD also impacts calcium buffering and export mechanisms.

### Kidney calcium buffering and circulating calcium is impacted by genetic variation

Because genetic variation contributes to SCD severity,^36^ we sought to identify genetic variants driving kidney calcium binding protein abundance by analyzing previously published protein quantitative trait loci (pQTL) data generated from kidneys of 6–18month-old diversity outbred (DO) mice (∼1:1 sex ratio).^37^ Each DO mouse contains a random assortment of genetic variants achieved by breeding together eight founder inbred strains: A/J; C57BL/6J; CAST/EiJ (CAST); 129S1/SvlmJ; NOD/ShiLtJ; NZO/H1LtJ; PWK/PhJ (PWK); and WSB/EiJ (WSB) (**Figure 5a**). Among the three main kidney proteins associated with calcium buffering in the distal tubule (CALB1, PVALB, S100G), the abundance of CALB1 and PVALB had strong pQTLs located in *cis* to their encoding genes on Chr 4 and Chr 15, respectively, overlapping the location of the genes’ eQTLs (**Figure 5b** and **Supplementary Figure 5a**). The eQTL and pQTL peaks of *Calb1* (chr4:14,855,973 and chr4:15,213,166 respectively) were located upstream of the *Calb1* gene body (**Figure 5c**). eQTL and pQTL profiles at the *Calb1* locus revealed a high correlation between them (r = 0.94) (**Figure 5d**), suggesting that levels of CALB1 protein and its encoding mRNA are controlled by the same underlying genetic variants. Founder strain allelic contributions derived from the eQTL and pQTL mapping model for the peaks on Chr 4 suggest that CAST has the lowest levels of CALB1 and its encoding mRNA (**Figure 5e**). Founder allele-effect estimates at the pQTL peak showed nearly identical direction and magnitude between the mRNA and protein profiles (r = 0.98) (**Figure 5f**).

**Figure 5:**
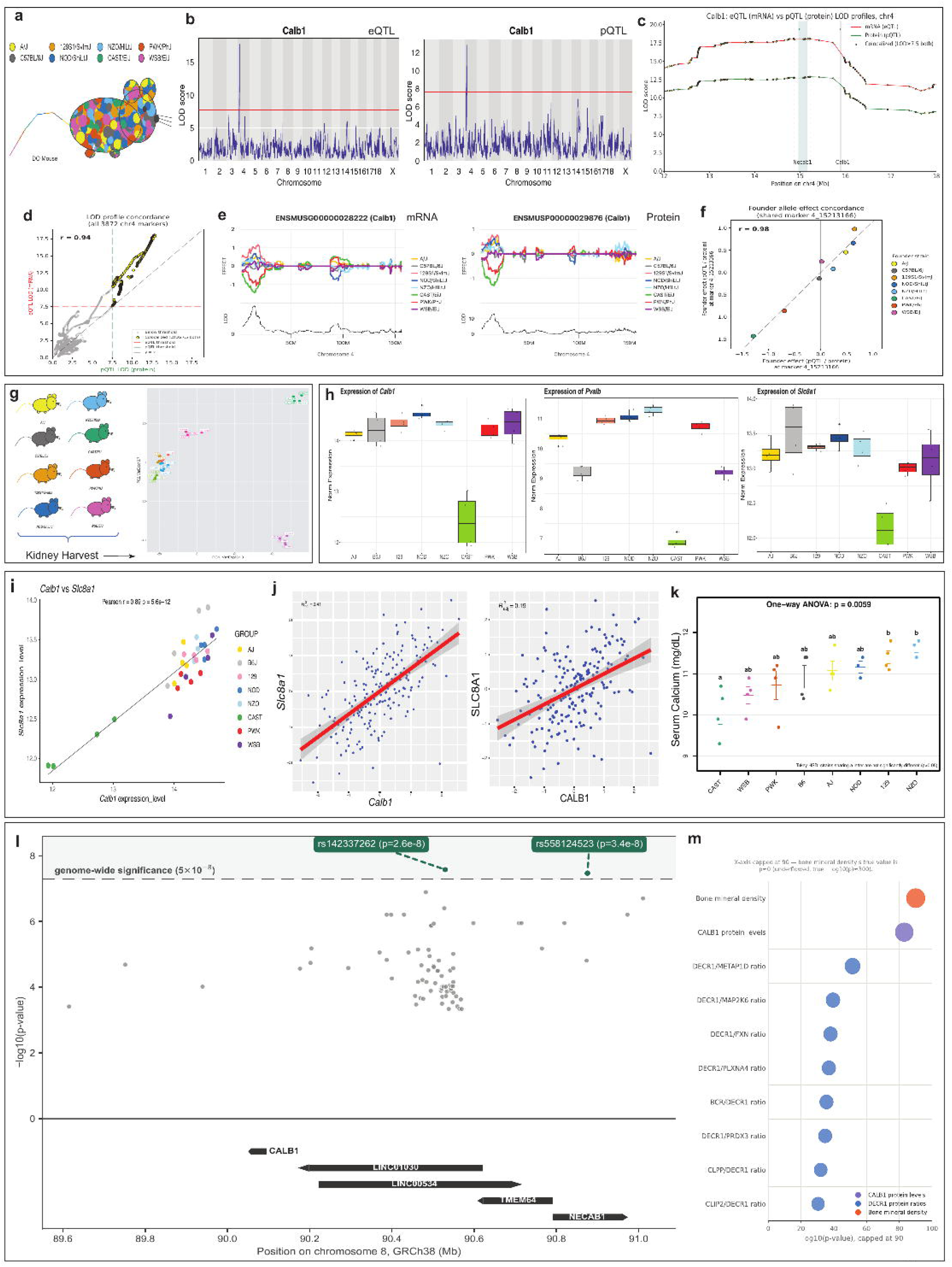
Levels of calcium buffers are impacted by genetic variation. (**a**) The Diversity Outbred mouse genome as a mosaic. Each Diversity Outbred (DO) mouse is a genetic mosaic of the same 8 inbred founder strains (A/J, C57BL/6J, 129S1/SvImJ, NOD/ShiLtJ, NZO/HlLtJ, CAST/EiJ, PWK/PhJ, WSB/EiJ, colored per the standard JAX founder palette), illustrated as a patchwork of founder colors across a single mouse silhouette. (**b**) Genome-wide LOD scores of kidneys from a 6-month-old DO mouse population^37^ reveals protein quantitative trait loci (pQTL) associated CALB1 abundance and overlapping expression QTL (eQTL) for its encoding gene in cis on Chr 4. (**c**) Colocalized eQTL and pQTL signals at the Calb1 locus. Genome-scan LOD profiles across chromosome 4 (12–18 Mb window) for Calb1 mRNA abundance (eQTL, red) and protein abundance (pQTL, green) in a DO founder-haplotype mapping panel. Yellow markers indicate positions where both scans exceed the significance threshold of LOD ≥ 7.5 (dashed gray line), showing that the two traits map to the same genomic interval. Gray- and teal-shaded regions mark the Calb1 and Necab1 gene bodies, respectively, with arrows indicating strand direction (Calb1, + strand; Necab1, − strand). Both QTL peaks fall just outside these gene bodies, within the intergenic region separating them. (**d**) Concordance between eQTL and pQTL signals at Calb1. (Left) LOD scores for the protein (pQTL) vs. mRNA (eQTL) genome scans at every chr4 marker; yellow points exceed the LOD ≥ 7.5 threshold in both scans. Most markers cluster along the diagonal (r = 0.94). (**e**) Founder allele effects plots revealed that the Calb1 eQTL and pQTL peaks on Chr 4 are driven by lower expression from the CAST/EiJ allele (green line). (**f**) Founder allele effects at the shared peak marker (4_15213166), comparing each of the eight DO/CC founder strains’ effect on protein vs. mRNA abundance. Strong agreement in direction and magnitude (r = 0.98) supports a single, shared causal variant driving both traits. (**g**) Kidneys were collected from eight DO founders followed by bulk RNA-seq. PCA plots based on mRNA levels revealed distinct strain-based clustering. (**h**) Kidney expression levels of Calb1, Pvalb, and Slc8a1 across DO founders. (**i**) Correlation between Calb1 and Slc8a1 expression using a kidney RNA-seq dataset generated from DO founder female mice. (**j**) Correlation between Calb1 and Slc8a1 expression using a kidney RNA-seq and proteomics dataset generated from DO male and female mice. (**k**) Serum calcium levels in DO female founders. (**l**) Regional association plot of cis-eQTL variants for CALB1 in kidney tissue. Each point represents one of 78 variants tested for association with CALB1 expression in a kidney dataset (N=474), plotted by genomic position (chr8, GRCh38) against −log10(p-value). Variants rs142337262 and rs558124523 exceed the genome-wide significance threshold (p<5×10⁻⁸, dashed line). Gene tracks below the plot show, to scale, the five annotated genes spanning this ∼1.5 Mb region: CALB1, the overlapping long non-coding RNAs LINC01030 and LINC00534, TMEM64, and NECAB1 (a second calcium-binding protein gene). (**m**) Top 10 GWAS Catalog trait associations at the CALB1 locus, ranked by significance. Bubble size and horizontal position both scale with −log10(p-value) (x-axis capped at 90 for readability). Bubble colors: CALB1 protein levels (purple), DECR1 protein-level ratios (blue), and bone mineral density (orange). The bone mineral density association (rs117035011) reflects a p-value that underflowed to zero in the source data (true −log10(p) ≈ 300, off the plotted scale).

To validate the predictive strain-dependent finding and directly assess gene expression variation across the eight DO founder strains, we profiled their kidney transcriptomes. The transcriptomes were highly variable among the strains, with PCA supporting ‘strain’ as a major factor driving variation. The wild-derived founder strains CAST, PWK, and WSB were most distinct, underlying much of the variability in gene expression in the kidney (**Figure 5g**). By testing the distribution of gene expression levels across the strains, we found that CAST had the lower end of *Calb1, Pvalb,* and *Slc8a1* expression when compared to the other strains (**Figure 5h**). Further, *Calb1* and *Slc8a1* expression in the kidneys of DO mice was correlated with the CAST allele associated with the lowest *Calb1* expression (**Figure 5i**), consistent with a strong genetic contribution to *Calb1* expression level variance. To determine whether the correlative trait of low *Calb1* and *Slc8a1* expression was inherited from the CAST founder, we analyzed mRNA and protein data from each DO founder strain. We found that *Slc8a1* mRNA and SLC8A1 protein were the top markers that correlated with levels of *Calb1* and CALB1, respectively, across the DO founders (**Figure 5j**). To determine whether genetic background associated with differential CALB1 and SLC8A1 abundance can contribute to circulating calcium levels, we measured serum calcium across the eight DO founders. CAST mice had low circulating calcium levels, suggesting that genetic variants in this strain may contribute to the circulating calcium heterogeneity (**Figure 5k**).

Similar to genetic variation at *Calb1* in mice that alters its renal expression level, our analysis of human kidney datasets^27^ derived from 474 individuals yielded 78 variants associated with levels of *CALB1* expression. Of these, two reached genome-wide significance (p<5×10⁻⁸) for *cis*-regulatory association with *CALB1* expression in kidney tissue: rs142337262 (chr8:90,529,441; p=2.62×10⁻⁸) and rs558124523 (chr8:90,875,410; p=3.38×10⁻⁸) (**Figure 5l**). The lead variant rs142337262 is located between the *CALB1* and *NECAB1* gene bodies. Follow-up annotation showed this variant directly overlaps introns of two long non-coding RNAs (*LINC01030, LINC00534*). Particularly, *LINC01030* genetic variants are associated with CALB1 protein levels and bone mineral density (**Figure 5m**). Taken together, these findings demonstrate that renal expression of calcium buffer genes and the abundance of their encoded proteins are significantly impacted by genetic variation, a driver of variable hypocalcemia status in SCD.

### CALB1 sustains mitochondrial oxidative capacity and FGF23 bioactivity via 2,3-disphosphoglycerate

Next, we tested if cellular changes in calcium binding alter energy metabolism in DCT cells. When comparing HbSS vs. HbAA kidneys for changes in the abundance of metabolic proteins (**Figure 6a**) using mass spectrometry, we found that glycolytic/energy-sensing proteins are predominantly upregulated while oxidative phosphorylation and tricarboxylic acid (TCA) cycle subunits are predominantly downregulated (Benjamini-Hochberg procedure -FDR q<0.05), consistent with a metabolic shift from mitochondrial respiration to glycolysis in SCD (**Figure 6b**).

**Figure 6:**
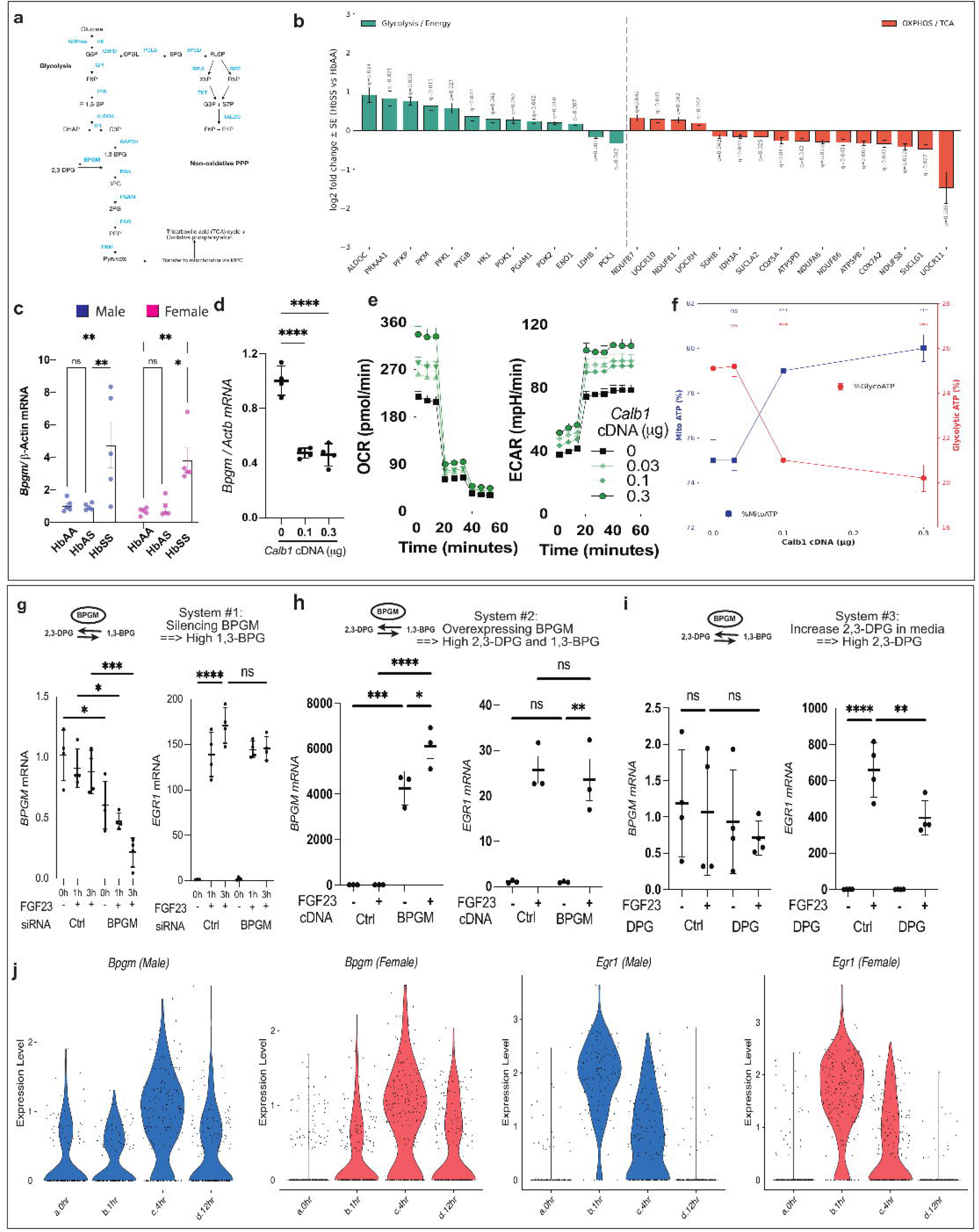
Calb1 rewires DCT cell energy metabolism. (**a**) Diagram of cellular energy pathways: glycolysis and pentose phosphate pathway (PPP) (metabolites, black; enzymes, blue) in the cytosol and the citric acid cycle and oxidative phosphorylation in the mitochondria. (**b**) Glycolysis and oxidative phosphorylation/tricarboxylic acid cycle protein abundances in kidneys of HbSS vs. HbAA mice using mass spectrometric analysis. (**c**) Bpgm mRNA levels in male and female HbAA, HbAS, and HbSS kidneys determined by RT-qPCR analysis. (**d**) DCT cell line 209 was transfected with Calb1 cDNA for 32 h followed by 16 h in hypoxic (1% O_2_) conditions and then subjected to RT-qPCR analysis of Bpgm. (**e**) Oxygen consumption rate (OCR), extracellular acidification rate (ECAR), total ATP production rate, and the percentage of glycolytic and mitochondrial ATP were measured under normoxic (21% O_2_) conditions by Seahorse assay in DCT 209 cells transfected with different amounts of Calb1 cDNA. (**f**) Calb1 cDNA dose-dependently increased mitochondrial ATP (XF Real-Time ATP Rate Assay, oligomycin + rotenone/antimycin A; mean ± SEM, n = 8/dose). (**g–i**) EGR1 and BPGM mRNA levels determined by RT-qPCR for mKL-HEK293 cells treated with BPGM siRNA (**g**), BPGM-encoding cDNA (**h**), or 2,3-DPG (**i**) followed by treatment with FGF23. (**j**) Violin plots of Bpgm and Egr1 expression in C57BL/6J male and female mice treated with FGF23 for 0, 1, 4, and 12 h. Plots were generated by analyzing our previous scRNA-seq dataset.^29^

To account for this apparent disconnect between increased glycolysis and mitochondrial energy production, we hypothesized that excess glycolytic flux triggers the Rapoport-Luebering shunt, a pathway that diverts 1,3-bisphosphoglycerate (BPG) from glycolysis by converting it to 2,3-DPG (**Figure 6a**). This conversion is catalyzed by BPG mutase (BPGM) encoded by *Bpgm,*^38^ whose expression was upregulated across different segments of the nephron in HbSS vs. HbAS mice (**Supplementary Figure** 6a). We confirmed this upregulation of *Bpgm* in kidneys of male and female HbSS vs. HbAS/HbAA mice (**Figure** 6c). To test whether calcium buffering potential influences the Rapoport–Luebering shunt, we transfected DCT cell line 209 with the *Calb1* cDNA and assessed the expression of *Bpgm*. We observed that *Calb1* overexpression decreased *Bpgm* expression (**Figure 6d**), consistent with the possibility that CALB1 levels contribute to metabolic reprogramming of DCT cells via the Rapoport–Luebering shunt.

To determine how calcium buffering potential affects energy metabolism in the DCT, we measured oxygen consumption rate (OCR) and extracellular acidification rate (ECAR) of DCT cell line 209 overexpressing *Calb1*. Using the Seahorse assay, we observed increased oxidative phosphorylation as shown by real-time OCR and ECAR profiles, and this effect was dependent on cDNA dose (**Figure 6e; Supplementary Figure 6b**). Moreover, analysis of cytosolic vs. mitochondrial ATP production revealed that *Calb1* overexpression increased the output of ATP production from mitochondria (**Figure 6f**).

To test if the Rapoport–Luebering shunt affects renal FGF23 signaling, we treated our previously generated human embryonic kidney cell line that produces membrane-bound KL (mKL-HEK293)^29^ with either *BPGM* siRNA, *BPGM*-encoding cDNA, or 2,3-DPG followed by addition of FGF23 and then measured expression of the canonical FGF23 signaling target, *EGR1*.^39^ Silencing *BPGM* which impedes the conversion of 1,3-BPG into 2,3-DPG^40^ did not change *EGR1* expression, suggesting that 2,3-DPG rather than 1,3-BPG influences FGF23 signaling (**Figure** 6g). Overexpressing *BPGM*, which has the potential to increase the amount of 1,3-BPG and 2,3-DPG,^41^ did not change *EGR1*, consistent with 2,3-DPG influencing FGF23 signaling (**Figure** 6h). Indeed, there was a decrease in *EGR1* when comparing 2,3-DPG + FGF23 to FGF23 alone (**Figure** 6i), suggesting that BPGM activity increases the amount of 2,3-DPG to influence FGF23 bioactivity. Finally, we tested whether FGF23 signaling regulates *Bpgm in vivo* by analyzing kidney scRNA-seq dataset derived from mice treated with FGF23.^29^ In DCT cells, *Egr1* expression rose 1 h after FGF23 administration in male and female mice (**Figure** 6j) and then decreased at 4 h before resetting to baseline at 12 h. *Bpgm* increased in DCT cells of male and female mice in response to FGF23, with a peak at 4 h after administration (**Figure** 6j). Taken together, our findings indicate that *Calb1* reprograms DCT cell glycolytic fate and BPGM signaling via 2,3-DPG, thereby fine tuning FGF23 signaling as a critical mechanism that controls calcium reabsorption in the DCT.

In summary (**Figure 7**), genetic and environmental factors like dietary iron intake that influence iron metabolism and hormonally mediated FGF23 signaling control calcium buffering potential in kidney DCT cells. In human and mouse, CALB1 and PVALB expression is influenced by genetic variation. We revealed that *CALB1* expression is controlled in a KL-dependent FGF23 signaling manner and that CALB1 controls DCT metabolism by rewiring BPGM activities. In a pathological state where KL is decreased, such as SCD, acute kidney injury, CKD and/or iron deficiency, levels of the calcium buffers CALB1 and PVALB are reduced. Our findings provide the first evidence that connects, at single-cell resolution, the iron and cellular metabolism defects in SCD with mineral metabolism alteration in kidney DCT cells.

**Figure 7:**
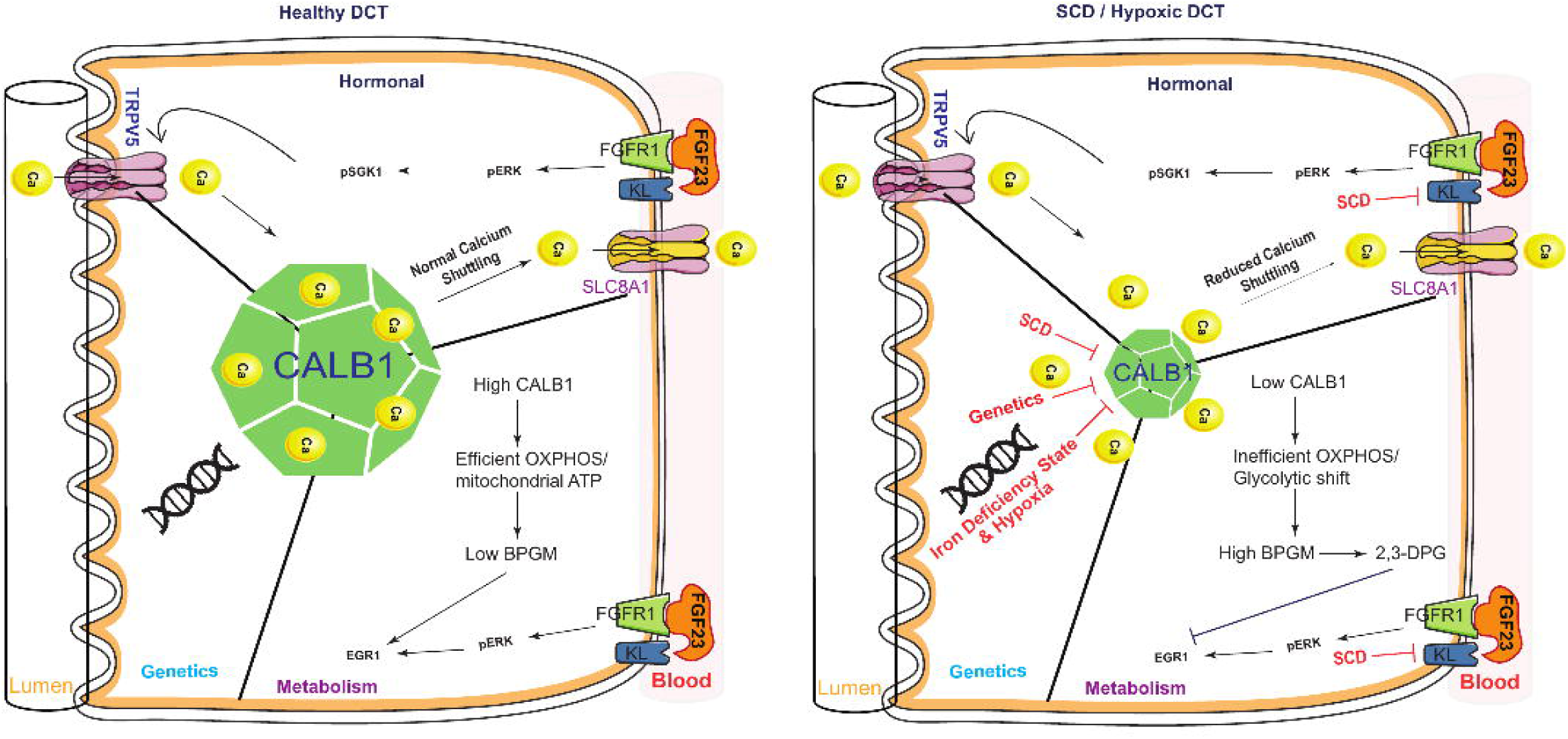
CALB1 expression, shaped by genetic variation and iron status, links renal calcium buffering to DCT energy metabolism and FGF23 bioactivity. In a healthy state, DCT cells produce a high level of CALB1 to sequester cytosolic calcium and contribute to its shuttling for export through SLC8A1. This mechanism is also essential to keep DCT cellular metabolism in an energy-efficient state. However, in SCD, hypoxia, or due to specific genetic mutation, CALB1 levels decrease in DCT cells, leading to altered metabolism and reduced shuttling and export of calcium. Changes in cellular metabolism designed for accumulation of 2,3-DPG as seen in SCD also trigger a feed-forward-loop mechanism that reprograms the efficiency of FGF23 signaling, a canonical regulator of renal calcium reabsorption.

## Discussion

Circulating calcium levels are heterogeneous in SCD. A previous prospective observational study revealed that most SCD patients exhibit hypocalcemia,^7^ the most common mineral bone abnormality seen in SCD patients, associated with altered VitD and PTH levels, whereas other SCD patients have relatively normal serum calcium levels.^7^ The reasons for this phenotypic heterogeneity have been unclear. The high heterogeneity of circulating calcium observed in our meta-analysis supports that genetic and environmental factors rather than a uniform disease mechanism drive variable calcium handling in SCD. Using the Townes mouse model of SCD, we showed that SCD mice have decreased serum calcium levels. We revealed that mechanisms associated with calcium reabsorption and trafficking in kidney DCT cells are altered in the SCD state and are modulated by genetic and environmental factors in addition to being hormonally controlled.

We found enrichment of the binding-site motif for the hypoxia-responsive transcriptional repressor REST^26^ in chromatin accessibility peaks at the *Calb1* locus in DCT cells in HbSS vs. HbAS mice, suggesting increased potential for REST binding in the SCD state. Hypoxia, a common SCD mechanism, increases intracellular calcium by altering the cellular energy supply that affects calcium channels and further increases intracellular calcium levels.^42^ It is possible that hypoxia could directly change the binding affinity of REST to RE1 sites in DCT, as previously proposed in a study of mouse hippocampal neural progenitor cells.^26^ In the nervous system, REST has been shown to repress neuronal gene transcription via RE1 elements.^43^ Based on our motif-enrichment analysis suggests, we speculate an analogous RE1-mediated repressive mechanism may operate at the *Calb1* locus in DCT cells. This could involve a REST co-repressor that remains to be identified in the kidney. Whether transcriptional regulation of *Calb1* in DCT cells mirrors that occurring in neurons is unknown. We determined that REST-mediated transcriptional downregulation of *Calb1* is controlled by KL-dependent FGF23 signaling. While the effects of PTH and VitD in regulating *Calb1* in the DCT remain to be fully determined, we found that neither circulating PTH levels nor the expression of *Pth1r* in DCT cells were increased in HbSS vs. HbAS mice. Overall, our data suggest a more prominent role for altered KL-dependent FGF23 signaling in reprogramming DCT calcium reabsorption capacity in SCD.

Our work in Townes mice also showed that SCD decreases CALB1 levels in the kidney, concomitant with low blood calcium. In accord with previous studies of neurons in *Calb1* knockout mice showing increased intracellular calcium due to impaired neuronal calcium buffering,^44^ the decrease of CALB1 in the kidney may increase intracellular calcium levels. Moreover, we revealed that misregulation of calcium binding potential changes distal tubule cellular metabolism. Increased glycolysis, particularly in the form of aerobic glycolysis, is observed in kidney diseases, and it was reported that glycolysis contributes to kidney disease progression.^45^ This suggests the possibility that therapeutic targeting of glycolysis could slow or reverse kidney damage^46^ and improve kidney function in SCD. Further preclinical studies could determine whether targeting excessive, yet energy-deficient, glycolysis in SCD improves kidney function. It is also possible that the excessive glycolysis observed in SCD results from glycolytic activity that is not effectively translated into mitochondrial energy output. Our findings are consistent with the possibility that BPGM-mediated conversion of the glycolytic metabolite 1,3-BPG into 2,3-DPG is increased in SCD, a mechanism that we expect would further limit energy production in DCT cells despite their excessive glycolysis.

Previous studies have demonstrated that increased activity of the Rapoport–Luebering shunt (i.e., BPGM/2,3-DPG pathway) suppresses glycolysis in hypoxic astrocytes.^47^ Altered glycolysis results in increased levels of 2,3-DPG, which promotes hemoglobin S polymerization and red blood cell (RBC) sickling, and reduced ATP production, thereby affecting RBC deformability and survival.^48,49^ It is possible that metabolic reprogramming similar to that observed in RBCs during SCD also occurs in kidney DCT cells. An intriguing translational implication of our findings concerns pyruvate kinase activators currently in clinical trials for SCD.^50,51^ Mitapivat and the related compound GBT021601 reduce intracellular 2,3-DPG in RBCs by activating pyruvate kinase, thereby shifting glycolytic flux away from the Rapoport–Luebering shunt and toward pyruvate production, which increases hemoglobin oxygen affinity and reduces sickling. Our data now suggest that 2,3-DPG is not merely an RBC-restricted regulator of hemoglobin activity but may also act in kidney distal tubule cells to limit KL-dependent FGF23 signaling, which is critical in calcium reabsorption. If mitapivat can limit the amount of 2,3-DPG in DCT cells in SCD patients, then it may exert a secondary, previously unrecognized benefit in the kidney by reducing 2,3-DPG levels and thereby restoring FGF23 signaling activity to improve calcium reabsorption. Conversely, any therapy that inadvertently raises renal 2,3-DPG could compound the renal calcium buffering defect in the SCD state. These possibilities can be tested clinically by monitoring serum and urinary calcium, as well as FGF23 levels, in SCD patients enrolled in ongoing mitapivat trials. More broadly, our findings suggest that the therapeutic benefit of reducing 2,3-DPG in SCD should be evaluated not only through the lens of hemoglobin oxygen affinity but also in the context of organ-level metabolic reprogramming, where this metabolite may unexpectedly reduce FGF23 signaling activity below a threshold required for effective calcium reabsorption.

Similar to the heterogeneity of serum calcium levels in SCD patients, their iron status in is highly variable,^52^ with patients frequently showing iron deficiency, sufficiency, or overload, often due to transfusion, inflammation, or a combination of factors. We reveal the role of dietary iron in reprogramming calcium reabsorption and trafficking in the kidney during SCD. Our study pinpoints how genetic architecture could predispose individuals to different levels of calcium buffering potential. Building on the instrumental contributions SCD mouse models^53,54^ have made to understanding SCD pathology, our study herein demonstrates the potential of genetic variation to influence SCD symptom severity at either the systemic level (blood calcium) and/or at the local level (calcium trafficking in the kidney DCT) with kidney CALB1 and PVALB levels being highly heritable and their associated *cis*-QTLs representing candidate genetic drivers of blood calcium heterogeneity. As we previously proposed,^55^ increasing the genetic diversity of SCD mouse models such as Townes and Berkeley could uncover variants that contribute to hypocalcemia in SCD and thereby lead to new therapeutic targets for SCD as well as open paths for personalized SCD therapeutic management in patient. We note that *CALB1* has not yet appeared in human SCD GWAS modifier screens, likely because those studies have been powered for fetal Hb-related and hematologic outcomes rather than renal mineral phenotypes. Future GWAS using serum calcium or urinary calcium as the outcome in SCD cohorts may identify *CALB1*-region variants as novel modifiers.

In sum, SCD disrupts calcium homeostasis by impairing renal calcium reabsorption pathways in the distal segments of the nephron. Collectively, the novel FGF23 signaling dysfunction in SCD and the potential for genetic predisposition as well as dietary factors to control renal calcium buffering provide new junction points for interventions in mineral metabolism that occur in SCD, CKD, and acute kidney injury.

## Materials and Methods

### Meta-analysis of human study data

PubMed was searched in July 2026 using combinations of “serum calcium,” “sickle cell disease/anemia,” and related terms (e.g., “hypocalcemia,” “vaso-occlusive crisis,” “calcium magnesium phosphorus”), with no date, language, or design restrictions. Serum calcium values for cases and controls were extracted directly from published studies (mean ± SD and sample size per group). For each study, a between-group standardized mean difference (Hedges’ g, with small-sample correction) and its standard error were calculated from the reported means and SDs, and these six effect sizes were pooled using a random-effects (DerSimonian–Laird) model, with between-study heterogeneity quantified via Cochran’s Q and I².

### Ethics statement for mouse studies and general mouse husbandry

All mouse studies were performed following approval by The Jackson Laboratory Institutional Animal Care and Use Committee (IACUC). All mice were housed in The Jackson Laboratory Research Animal Facility (Bar Harbor, Maine) and fed a standard 6% 5KOG diet unless otherwise indicated. For tissue and blood collection, mice were euthanized with carbon dioxide.

### Mice

Wild-type (HbAA), heterozygous (HbAS), and homozygous (HbSS) Townes mice (B6;129-*Hbb^tm2(HBG1,HBB*)Tow^*/*Hbb^tm3(HBG1,HBB)Tow^ Hba^tm1(HBA)Tow^*/J were obtained from The Jackson Laboratory [#013071]). Mice carrying the doxycycline-inducible Dmp1-Tet3G/TRE3G-hFGF23(R176Q/R179Q)-IRES-hrGFPII transgene (FGF23-Tg) were obtained from Dr. Kenneth E. White at Indiana University and used as previously described^30^. Briefly, osteocyte-restricted overexpression of human *FGF23(R176Q/R179Q)* was induced in transgene heterozygotes by administering 2 mg/ml of doxycycline in the drinking water for 2 weeks followed by euthanasia. DO founder mice were obtained from The Jackson Laboratory (Bar Harbor, Maine) and euthanized at 12–14 weeks of age.

### Mouse diet treatments

For iron deficiency, 5-week-old mice were randomized to receive either an iron-replete control diet (TD.09772, formulated with 45 ppm added Fe as ferrous sulfate heptahydrate, plus a background iron content of 2–6 ppm) or an iron-deficient diet (TD.09771, background iron content 2–6 ppm, no supplemental iron; Inotiv, Madison, WI) for four weeks, after which mice were euthanized. For adenine supplementation, 8-week-old mice were put on a normal casein diet (Inotiv, TD.200790) or adenine diet (0.2%) (Inotiv, TD.200791) for two or four weeks and then euthanized.

### Spleen imaging

A cohort of male HbSS, HbAS, and HbAA mice were intravenously administered a single dose of the ExiTron™ nano12000 contrast agent and imaged live using the Quantum GX micro-CT system at 5, 6, and 8 weeks of age. Analyze 12.0 software was used to derive total spleen volumes from their cross-sectional areas in segmented images.

### Multiome single-nuclei RNA-seq and single-nuclei ATAC-seq

Whole male and female kidney pairs were finely minced and the tissue dissociated using a chemical (detergent-based) lysis process containing Nonidet P40 to generate a suspension of single nuclei. For each sample, 10,000 cell recovery was targeted and applied a single cell master mix with lysis buffer and reverse transcription reagents, according to the Chromium Single Cell 3’ Reagent Kits v3 User Guide, CG000183 Rev A (10x Genomics) for combined single nuclei RNAseq and ATACseq. Briefly, following nuclear dissociation, single-nucleus multiomics libraries were generated using Chromium Next GEM Single Cell Multiome ATAC+ Gene Expression Reagent Kit (10× Genomics).Multiome snRNA-seq and snATAC-seq libraries were processed with Cell Ranger ARC v2.0.2 against the refdata-cellranger-arc-mm10-2020-A-2.0.0 reference. Downstream analysis used Seurat v4.2.0 and Signac v1.8.0 under R 4.2.1, with gene annotation from EnsDb.Mmusculus.v79 (mm10). Ambient RNA contamination was estimated and removed with SoupX v1.6.1, and doublets were removed with DoubletFinder v2.0.3. Gene expression was normalized with SCTransform and, in parallel, log-normalized (scale factor 10,000) with the 3,000 most variable features selected by the variance-stabilizing transformation; in both cases mitochondrial content, ribosomal content and RNA library size were regressed out. Chromatin accessibility was normalized by term frequency–inverse document frequency, variable features were selected with FindTopFeatures (min.cutoff = 20), and dimensions were reduced by singular value decomposition (latent semantic indexing, LSI). The two modalities were integrated by weighted nearest-neighbor analysis using Harmony-corrected dimensions 1–15 for RNA and 2–15 for ATAC. Clusters were called on the weighted nearest-neighbor graph, and UMAP and t-SNE embeddings were computed from the same graph. Clusters were annotated into seven populations using canonical markers of kidney cells. The sex of each nucleus was inferred from sex-linked transcripts: nuclei with normalized expression of any Y-linked gene (*Kdm5d, Eif2s3y, Uty, Ddx3y*) of at least 0.1 were assigned male, nuclei with *Xist* of at least 1 or *Tsix* of at least 0.5 were assigned female. Differential chromatin accessibility was tested separately within each annotated cell type. Peaks were compared between the iron-overload and iron-replete conditions with FindMarkers on the peak assay using logistic regression (test.use = “LR”) with the number of peak counts per nucleus as a latent variable to control for library complexity. Peaks were tested when detected in at least 10% of nuclei in either condition (min.pct = 0.1) with an absolute log_2_ fold-change of at least 0.25, requiring at least 20 nuclei per feature and per group, and both increased and decreased accessibility were retained. Peaks were annotated to overlapping gene features using the EnsDb annotation. KEGG pathway analysis was performed using ShinyGO.^56^ Transcription factor binding-site motif analysis was performed using the JASPAR package in R.

### Publicly available datasets

1. FGF23-treated male and female C57BL6/J (B6) mice scRNA-seq datasets^29^ are at the Gene Expression Omnibus (GEO): GSE246385.
2. Mouse IRI kidney scRNA-seq datasets^34^ are at GEO: GSE139107. Related figures were generated using the data visualization tool at the Humphreys Lab Kidney Interactive Transcriptomics home page (http://humphreyslab.com/SingleCell/) applied to dataset: “Mouse IRI Kidney (Sham, 4hrs, 12hrs, 2days, 14days,and 6weeks): Kirita et al, PNAS 2020).”
3. Human kidney scRNA-seq dataset was from the Kidney Precision Medicine Project (KPMP) Consortium.^24^
4. Human kidney combined scRNA-seq + snATAC-seq datasets.^27^
5. DO mouse pQTL and eQTL kidney datasets.^37^

### Mouse QTL mapping

Kidney transcriptome, proteome, and genotype data from the JAX Shock Center Diversity Outbred (DO) kidney study (n = 188 mice) were obtained from the Churchill Lab QTL Viewer (https://churchilllab.jax.org/qtlviewer/JAC/DOKidney). Transcript and protein abundances were each rank-normal transformed and mapped identically against 8 founder haplotype probabilities with qtl2::scan1(), using an additive model with DO generation, sex, and age as fixed covariates and a polygenic random effect based on leave-one-chromosome-out kinship matrices. Genome-wide significance thresholds were derived from 1,000 permutations (qtl2::scan1perm()): LOD = 7.77 (α = 0.05) and 5.86 (α = 0.63, suggestive) for transcripts, and LOD = 7.68 and 5.83 for proteins. At each peak marker, founder allele effects were estimated with qtl2::fit1(). Loci were refined by association mapping across a ±1 Mb window using qtl2::scan1snps() with founder variants from the Collaborative Cross variant database (fv.2021.snps.db3).

### Human eQTL plot

*Cis*-eQTL summary statistics (variant identifier, position, effect allele, effect size, standard error, and p-value) for *CALB1* in kidney tissue were obtained from a kidney dataset (N=474) and used without re-analysis. Genomic coordinates and strand orientation for *CALB1* and four neighboring genes (*LINC01030*, *LINC00534*, *TMEM64*, *NECAB1*) were retrieved from Ensembl (GRCh38). Variants were plotted by genomic position against −log10(p-value), with genome-wide significance defined as the conventional threshold of p<5×10⁻⁸. Gene tracks were drawn to scale beneath the association plot using the retrieved coordinates, with arrows indicating strand direction.

### Cell culture and transfection

Unless otherwise indicated, all cultured cells were incubated at 37°C in 5% CO_2_ and ambient oxygen conditions of ∼18%–20% O_2_. Our membrane-bound Klotho (mKL)-expressing HEK293 cell line^29^ (mKL-HEK293) was cultured in Eagle’s Minimum Essential Medium (ATCC, 30-2003) supplemented with 10% fetal bovine serum (Corning, MT35016CV). 209/MDCT cells (ATCC, CRL-3250) were cultured in a 1:1 mixture of Dulbecco’s Modified Eagle’s Medium (DMEM) with 1 g/L glucose and 1 mM sodium pyruvate (Gibco, 11-765-054) and Ham’s F-12 Nutrient Mix (Gibco, 11-765-054) supplemented with 5% fetal bovine serum (Corning, MT35016CV) and 1% penicillin-streptomycin (Millipore Sigma, P4333-100ML). Cells were transfected for 48 hours using FuGENE 4K Transfection Reagent (FuGENE, 4K-1000) with a 3:1 ratio of μL of FuGENE to μg cDNA.

### DPG and FGF23 treatment in vitro

mKL-HEK293 cells were plated into a 24-well plate in complete medium at 300,000 cells/ well. After 5 h, cells were cultured for 16 h in 10 mM 2,3-DPG (Millipore Sigma, D5764; reconstituted in molecular grade water) or in 0 mM 2,3-DPG as a control. Subsets of these 2,3-DPG-treated and non-2,3-DPG-treated cells were then cultured for 1 h in media containing 50 ng/mL FGF23 (Bio-Techne, 2604-FG-025/CF; reconstituted in and serially diluted with 1x PBS), 10 mM 2,3-DPG + 50 ng/mL FGF23, or 1x PBS as a control. RNA was then extracted. Four replicates of this experiment were performed.

### BPGM siRNA and FGF23 treatment in vitro

mKL-HEK293 cells were plated into two 6-well plates in complete medium at 300,000 cells/well. After 24 h, cells were treated for three days with either BPGM siRNA (Horizon Discovery, L-008917-00-0005) or control siRNA (Horizon Discovery, D-001720-03-05) at 8 nM, or vehicle alone. siRNAs were reconstituted in 1x siRNA buffer (Horizon Discovery, B-002000-UB-100) at 50 µM prior to use. Subsets of the siRNA-treated cells and untreated control cells were then cultured for 1 h in media containing 50 ng/mL FGF23 (as described above). RNA was then extracted. This experiment was performed in triplicate.

### BPGM cDNA and FGF23 treatment in vitro

mKL-HEK293 cells were plated into two 6-well plates in complete medium at of 300,000 cells/well, with three wells per treatment condition. After 24 h, cells were transfected using Fugene (FuGENE, 4K-1000) with 1 µg of a BPGM-encoding, GFP-tagged ORF expression vector (OriGene, RG202105) or empty expression vector pCMV6 (OriGene, PS100001). After 1 day, a subset of the transfected cells and untransfected cells were cultured for 1 h in media containing 50 ng/mL FGF23 (as described above). RNA was then extracted. This experiment was performed in triplicate.

### RNA extraction from cultured cells

Following respective treatment periods, culture medium was aspirated and wells rinsed with 1 mL of 1x PBS. Three hundred thousand cells were then collected by scraping and resuspended in 1 mL of 1x PBS. For RNA extraction, 500 µL of cells per well were spun at 500 g for 10 min in a cold microfuge at 4°C and RNA extracted using the Quick-RNA Miniprep Kit (Zymo Research, R1055). RNA integrity and concentration were quantified using the Nanodrop Spectrophotometer for quality control.

### Reverse transcription–quantitative PCR (RT-qPCR) analysis

Samples of total RNA were converted to cDNA via using the TaqMan™ RNA-to-CT™ 1-Step Kit (Applied Biosystems/Thermo Fisher, 4392938) with 20ng of RNA as a starting material and tested using pre-optimized primers (Applied Biosystems/Thermo Fisher) targeting mRNAs of genes listed in **Supplementary Table S1**) with *ACTB* or *Actb* used as an internal control, depending on sample species. Real-time PCR was performed on a QuantStudio 7 Flex real-time system (Applied Biosystems/Thermo Fisher) using TaqMan One-Step RT-PCR kit. PCR conditions for all experiments were: 30 min at 48°C, 10 min at 95°C, followed by 40 cycles of 15 s at 95°C and 1 min at 60°C. mRNA levels were calculated using the method described by Livak and Schmittgen^57^ as fold change (2-ΔΔCt) relative to the housekeeping β-actin genes *ACTB*/*Actb* with appropriate controls.

### ELISA of FGF23 and PTH

FGF23 levels in mouse plasma were quantified by subjecting 5-fold dilutions of each sample (in 1x PBS) to the MICROVUE Mouse/Rat FGF-23 (Intavct) ELISA Kit (QuidelOrtho, 60-6800), per manufacturer’s instructions. Intact FGF23 measures only the active, full-length hormone, while C-terminal FGF23 measures both the active hormone and its inactive breakdown fragments. PTH levels in undiluted mouse plasma were quantified via MICROVUE Mouse PTH 1–84 ELISA Kit (QuidelOrtho, NC0195335), per manufacturer’s instructions.

### Immunoassay of kidney proteins

20 mg of kidney tissue were lysed in Cell Lysis Buffer (10X) (Cell Signaling, 9803) with AEBSF (4-(2-Aminoethyl) benzenesulfonyl fluoride hydrochloride) Protease Inhibitor (Fisher Scientific, 78431) and incubated for 5 min at 4°C. Lysates were spun 10 min at 14,000 g in a cold microfuge at 4°C using a pre-chilled rotor to isolate the supernatant. Protein concentrations were determined using Bradford Reagent (Millipore Sigma, B6916). Capillary immunoassays were performed using the ProteinSimple Capillary Cartridge kit (ProteinSimple/Bio-Techne, SM-W004) and antibodies against KL (Proteintech, 28100-1-AP) and β-actin (Cell Signaling, 4970) for protein separation and detection, per manufacturer’s instructions, and immunosignals detected on a Wes instrument (ProteinSimple/Bio-Techne).

### Proteomics sample preparation and data acquisition

#### Sample Preparation for Proteomics Analysis

Kidney samples were thawed and 2:2:1 methanol:acetonitrile:water extraction buffer was added at 65 µL per 1 mg of tissue. All samples had a pre-chilled 5-mm stainless steel bead (QIAGEN) added and were lysed using a Tissue Lyser II (QIAGEN) for 2 min at 30 Hz. Stainless steel beads were removed using a magnet and samples were then ice-waterbath sonicated at 37 Hz (100% power; Firsherbrand FB11207) for 5 min (30 s on, 30 s off). Extractions were then incubated overnight in the -20°C, followed by centrifugation at 21,000 g at 4 °C for 15 min to pellet the protein. Protein pellets were reconstituted in 250 µL of 50 mM HEPES with 6 M Urea by pipetting, then using a Tissue Lyser II (QIAGEN) for 2 min at 30 Hz, and waterbath sonication as in the previous step. Lysates were clarified by centrifugation at 2,000 g at 4 °C for 15 min to remove heavy debris; supernatant was transferred to a new tube. Each sample had a 1:50 dilution (sample:50 mM HEPES) created and a microBCA assay (Thermo) was run for protein quantification according to the manufacturer protocol.

For each sample lysate, 20 µg of protein was diluted in 50 mM HEPES, pH 8.2 to bring the volume to 50 µL (diluting the Urea to < 1 M) for the trypsin digest. Protein reduction, alkylation, tryptic digestion, and subsequent Millipore C18 zip-tip (Millipore, ZTC18S096) peptide purification was then carried out as previously described.^58,59^ All purified peptides were dried in a vacuum centrifuge, then reconstituted in 20 µL of 98% H2O/2% ACN with 0.1% formic acid by vortex for 30 s at max speed. Liquid was then brought to the bottom of the tube using a low-speed tabletop minicentrifuge and reconstituted peptides were transferred to a mass spec vial (Thermo). Prior to analyzing samples via tandem mass spectrometry analysis, the samples were placed in the Vanquish Neo (Thermo) liquid chromatography autosampler set to 4 °C.

#### Liquid Chromatography Tandem Mass Spectrometry (LC-MS/MS) Analysis

All peptide samples were analyzed using the Thermo Orbitrap Astral mass spectrometer coupled to the Vanquish Neo liquid chromatography system with an EASY-Spray column (PepMap Neo C18, 75 µm x 150 mm, #ES75150PN) in The Jackson Laboratory Mass Spectrometry and Protein Chemistry laboratory as described previously.^58^ Following sample randomization (random.org), data independent acquisition mass spectrometry analysis (DIA-MS/MS) was performed on two technical replicates with an 8 µL injection for all samples. The Thermo Vanquish Neo liquid chromatography 25 minute gradient utilized Buffer A (100% H2O with 0.1% formic acid) and Buffer B (100% acetonitrile with 0.1% formic acid) with a flow rate of 600 nL/min. The full gradient conditions were run exactly as described previously.^58^ All Thermo Orbitrap Astral mass spectrometry DIA-MS/MS proteomics analysis was also performed as previously described in detail.^58^

#### DIA-MS/MS Proteomics Data Analysis

The RAW data files (.raw) from the Orbitrap Astral were automatically processed using the Thermo Ardia Server coupled directly to a Proteome Discoverer 3.1 workstation. The .raw files for each sample were searched against the UniProtKB Mus musculus (sp_tr_incl_isoforms TaxID=10090; v2024-05-01) protein database via CHIMERYS with inferys 3.0 in Proteome Discoverer 3.1. Search parameters included trypsin digestion, one missed cleavage maximum, 7–30 amino acids peptide length, charge state from +1 to 4, a 10 ppm fragment mass tolerance, dynamic oxidation of methionine (+15.995 Da), static carbamidomethlyl cysteine alkylation (+57.021 Da), and a false discovery rate of less than 0.05. All other Proteome Discoverer parameters followed manufacturer default recommendations. Data across the file was normalized to the total ion signal in the samples and a protein abundance-based ratio calculation was used for the summed abundances, followed by a background based t-test according to Proteome Discoverer guidelines. All sample data then underwent multiple corrections testing via Benjami-Hochberg application for an adjusted p-value. Strict parsimony was applied for protein grouping and the data was run with and without low abundance resampling imputation for comparison during the analysis process.

### Seahorse assay

The mouse DCT cell line 209 was plated in a 6-well plate (Corning; Costar Cat. #3516) at 300,000 cells per well. After 16 h, cells were transfected with mouse *Calb1* cDNA (OriGene, MR203429) for 2 days, resuspended in cell culture media before the extracellular flux analysis using the Seahorse XFe96/XF Pro FluxPak Mini (Agilent Technologies, 103793-100) assay kit. Before use, sensor cartridges were loaded with Calibrant Solution and left overnight in a non-CO_2_ incubator to hydrate the probe tips. Seahorse cell culture microplates were coated overnight with poly-D-lysine (Sigma Aldritch, P0899-10MG) then washed three times with 1x PBS (Cytiva). On the morning of the assay, resuspended 209 cells were split and plated on the pretreated microplates at 1.0 x10^5^ cells per well in Seahorse XF DMEM (Agilent Technologies, 103575-100) and then incubated for 1 h at 37°C in a non-CO_2_ incubator to allow CO_2_ diffusion followed with the assay. After oligomycin and rotenone/antimycin A from the Seahorse XF Real-Time ATP Rate Assay Kit (Agilent Technologies, 103592-100) were loaded into the sensor cartridge, the cells were assayed on the Agilent XFe/XF96 using Wave Software to determine OCR, ECAR, ATP production rate, and the percentage of ATP production from the cytosol and mitochondria.

### Bulk RNA sequencing

Total kidney RNA was isolated from cell pellets using the NucleoMag RNA Kit (Macherey-Nagel) and the KingFisher Flex System (Thermo Fisher Scientific). Cells were homogenized in MR1 buffer (Macherey-Nagel) by vortexing. RNA isolation was performed according to the manufacturer’s protocol. RNA concentration and quality were assessed using the Nanodrop 8000 spectrophotometer (Thermo Scientific) and the RNA ScreenTape Assay (Agilent Technologies). For RNA library preparation and sequencing, strand-specific libraries were constructed using the mRNA Library Prep Kit (Watchmaker), according to the manufacturer’s protocol. Briefly, polyA-containing mRNA were isolated using oligo-dT magnetic beads followed by RNA fragmentation, first- and second-strand cDNA synthesis, ligation of Illumina-specific adapters containing a unique barcode sequence for each library, and PCR amplification. The quality and concentration of the libraries were assessed using the D5000 ScreenTape (Agilent Technologies) and Qubit dsDNA HS Assay (ThermoFisher), respectively, according to the manufacturers’ instructions. Libraries were subjected to 150-bp paired-end sequencing on an Illumina NovaSeq X Plus using the 10B Reagent Kit.

### Quantification and statistical analysis

Statistical analyses were performed using GraphPad Prism (v10) or R (v4.2.1). One-way ANOVA with Tukey’s or Dunnett’s post hoc test was used for single-factor comparisons (e.g., in vitro dose-response treatments, genotype comparisons within one sex and time point). Two-way ANOVA was used for factorial designs — genotype × diet (HbAA/HbSS × IronCtrl/IronDef), genotype × diet duration (CAS/AD × 2 or 4 weeks), and genotype × time post-FGF23 (0, 1, 4, 12 h) — with Tukey’s post hoc test where interaction effects were significant. Correlation coefficients were compared across time points using Fisher z-transformation. Proteomic differential abundance used a background-based t-test with Benjamini-Hochberg correction (FDR q<0.05). Means ± SD (or SEM, as indicated) are shown in bar graphs. Significance was set at P<0.05 (or FDR q<0.05 for proteomic/genomic analyses).

## Acknowledgments

We thank Dr. Gary A. Churchill (The Jackson Laboratory) for his expert guidance on the genetic mapping study design and for valuable scientific discussions throughout the course of this work. This work was supported by the National Institutes of Health under grant numbers R00-DK129705 and R35-GM160386. The content is solely the responsibility of the authors and does not necessarily represent the official views of the National Institutes of Health. We thank the scientific services cores of Mass Spectrometry and Proteomics, Genome Technologies, and Single Cell Biology at The Jackson Laboratory for outstanding support. We thank Dr. Carmen C. Robinett for assistance with the manuscript. Figure illustrations were created using Servier Medical Art (https://smart.servier.com/), licensed under CC BY 4.0 (https://creativecommons.org/licenses/by/4.0/).

## Disclosure

The authors have nothing to declare.

## Author’s contributions

R.A. designed the study and conceived the experiments; G.X., K.J., E.W., and R.A., performed the *in vivo* and *in vitro* experiments; R.A. conducted cell/nuclei preparation procedure for single-cell biology experiments; G.X., Y-T.W., and R.A., performed bioinformatics analysis and interpretated the data. R.A. drafted and wrote the manuscript. All the authors edited and approved the submitted content of the manuscript.

## Data availability

All raw sequencing data (single-nuclei RNA-seq, single-nuclei ATAC-seq, and bulk RNA-seq) generated for this study have been uploaded to NCBI’s Gene Expression Omnibus (GEO): GSE348497.

---

**Supplementary Figure 1.** (**a**) Quality control (QC) plots for Townes mice HbAS (normal state) and HbSS (SCD state) multiomic datasets. (**b**) Unsupervised uniform manifold approximation and projection (UMAP) clustering comparison of male and female mice with sex-specific markers. (**c**) Dot plot of KEGG pathway enrichment in PT-S1 of HbSS vs. HbAS. (**d**) Violin plots show expression of calcium buffer-encoding genes *Pvalb* and *S100g* across all cell types in the kidney and in HbAS and HbSS. (**e**) Feature plot of *PVALB* expression from human kidney scRNA-seq data, highlighting its localized expression in DCT cells.

**Supplementary Figure 2.** (**a**) Read coverage and chromatin accessibility of the *CALB1* locus in human kidney cell types profiled by snATAC-seq.^27^ (**b**) Transcription factor binding-site motif enrichment in HbSS vs. HbAS. (**c**) Feature plots of mouse and human *VDR, PTHR1, KL,* and *FGFR1*. (**d**) Violin plots show *Kl, Fgfr1, Pth1r,* and *Vdr* expression in different kidney cell types of HbAS vs. HbSS male and female mice. (**e**) VDR binding-site motif enrichment analysis in HbSS vs. HbAS. (**f**) Blood plasma PTH levels in female and male HbAS and HbSS mice measured by ELISA.

**Supplementary Figure 3.** (**a**) Blood urea nitrogen (BUN) measured by colorimetric assay. (**b**) PCA plot of bulk RNA-seq data from male and female CKD kidney samples. (**c**) UMAP clustering identified various renal cell populations. (**d**) *Havcr1* expression dot plot from publicly available acute kidney injury dataset.

**Supplementary Figure 4.** ATP2B4 protein abundance measured by mass spectrometric analysis.

**Supplementary Figure 5.** Genome-wide LOD scores of kidneys from a DO mouse population^37^ reveals protein quantitative trait loci (pQTL) associated PVALB abundance and overlapping expression QTL (eQTL) for its encoding gene in *cis* on Chr 15.

**Supplementary Figure 6.** (**a**) *Bpgm* expression across renal cell types in HbSS vs. HbAS from our scRNA-seq datasets. Dot size represents the percentage of cells within each cell type that express *Bpgm* (percent expressed), and color intensity represents the scaled average expression level (average expression scaled/normalized across cell types) of *Bpgm* in that cell type. (**b**) DCT cell line 209 was transfected with different amounts of *Calb1* cDNA for 48 h and the expression levels analyzed by RT-qPCR.

**Supplementary Table S1:**
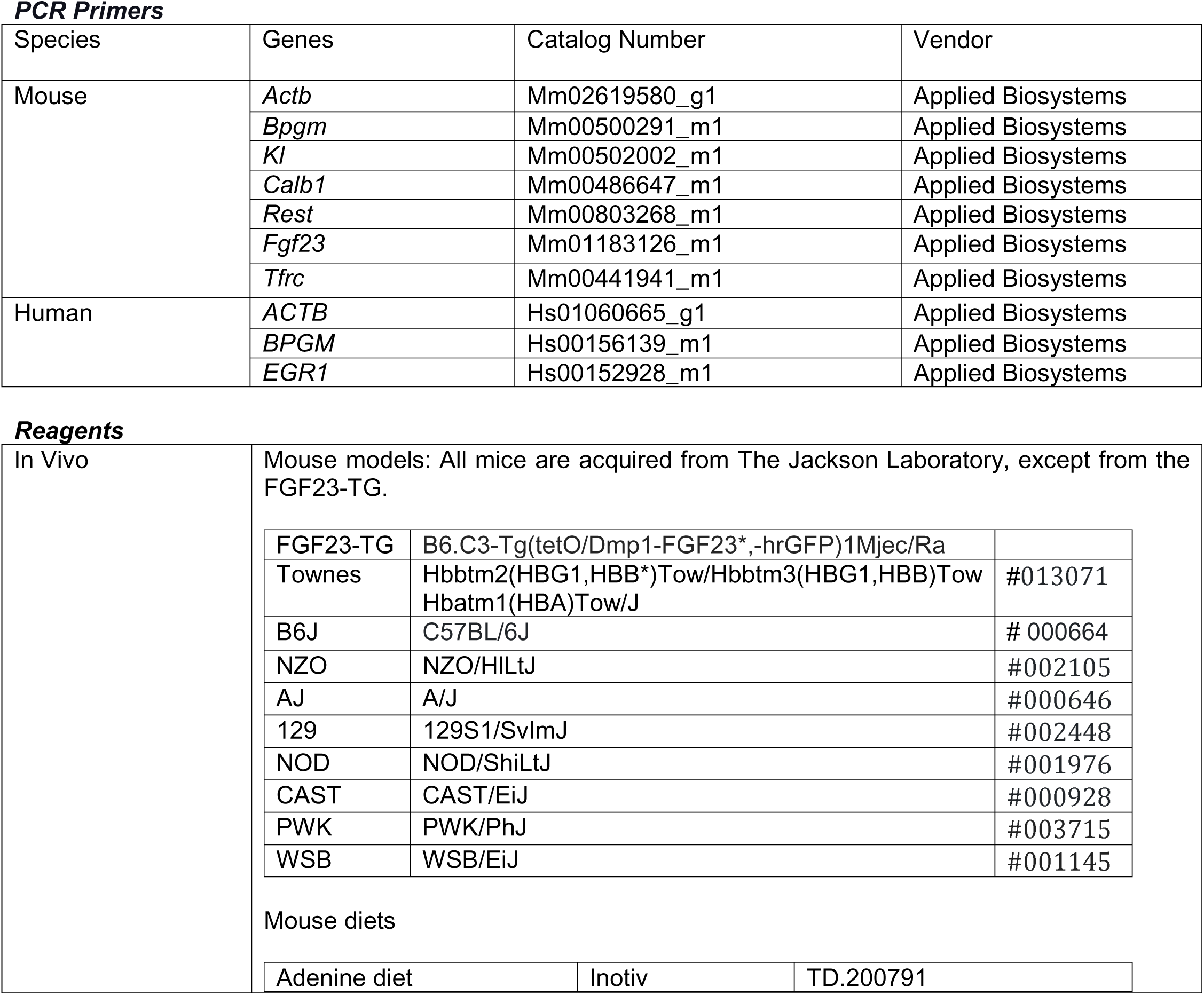
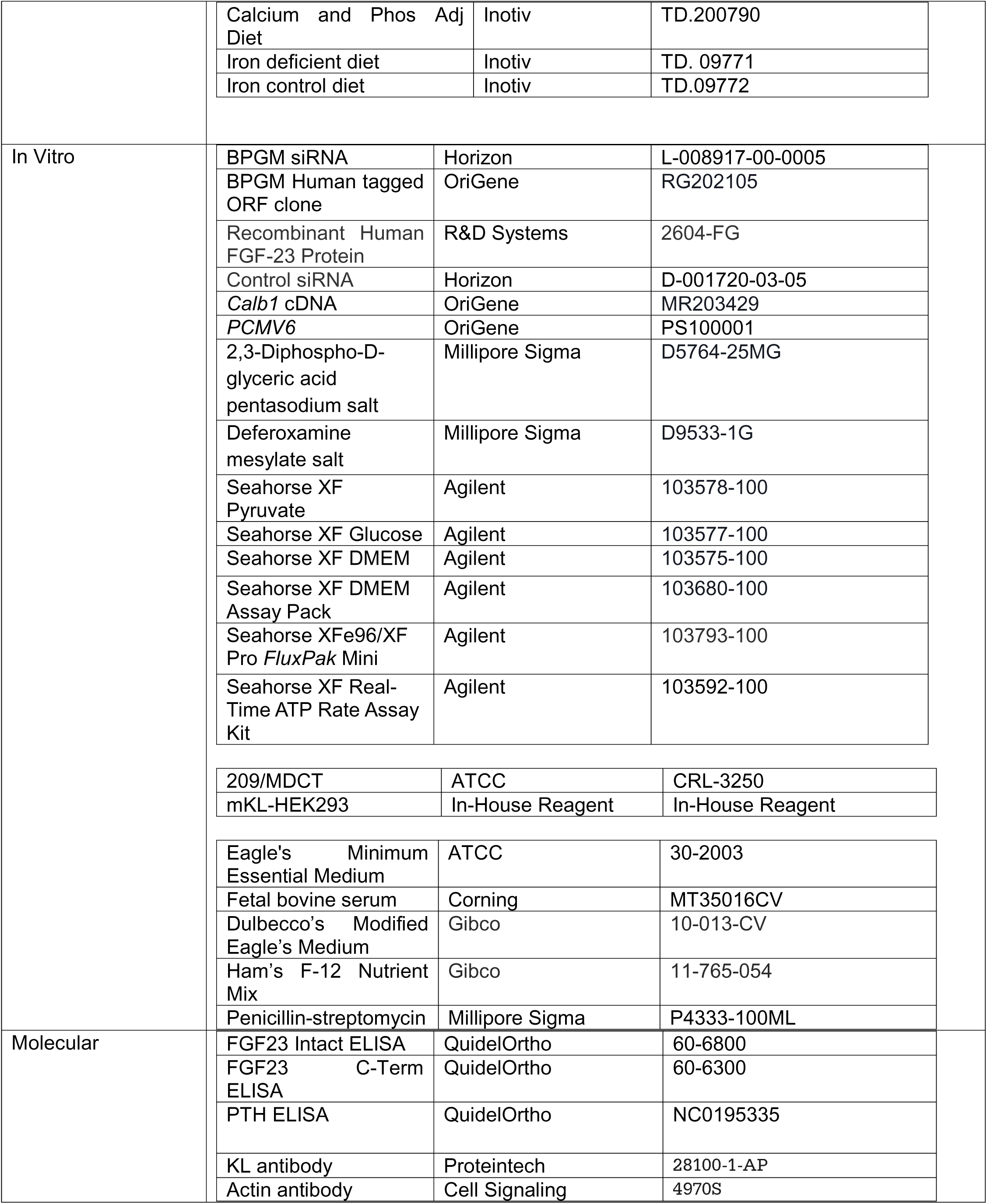

| ***PCR Primers*** |  |  |  |
|---|---|---|---|
| Species | Genes | Catalog Number | Vendor |
| Mouse | *Actb* | Mm02619580_g1 | Applied Biosystems |
|  | *Bpgm* | Mm00500291_m1 | Applied Biosystems |
|  | *Kl* | Mm00502002_m1 | Applied Biosystems |
|  | *Calb1* | Mm00486647_m1 | Applied Biosystems |
|  | *Rest* | Mm00803268_m1 | Applied Biosystems |
|  | *Fgf23* | Mm01183126_m1 | Applied Biosystems |
|  | *Tfrc* | Mm00441941_m1 | Applied Biosystems |
| Human | *ACTB* | Hs01060665_g1 | Applied Biosystems |
|  | *BPGM* | Hs00156139_m1 | Applied Biosystems |
|  | *EGR1* | Hs00152928_m1 | Applied Biosystems |
| ***Reagents*** |  |  |  |
| In Vivo | Mouse models: All mice are acquired from The Jackson Laboratory, except from the FGF23-TG. |  |  |
|  | FGF23-TG | B6.C3-Tg(tetO/Dmp1-FGF23*,-hrGFP)1Mjec/Ra |  |
|  | Townes | Hbbtm2(HBG1,HBB*)Tow/Hbbtm3(HBG1,HBB)Tow Hbatm1(HBA)Tow/J | #013071 |
|  | B6J | C57BL/6J | # 000664 |
|  | NZO | NZO/HlLtJ | #002105 |
|  | AJ | A/J | #000646 |
|  | 129 | 129S1/SvImJ | #002448 |
|  | NOD | NOD/ShiLtJ | #001976 |
|  | CAST | CAST/EiJ | #000928 |
|  | PWK | PWK/PhJ | #003715 |
|  | WSB | WSB/EiJ | #001145 |
|  | Mouse diets |  |  |
|  | Adenine diet | Inotiv | TD.200791 |
|  | Calcium and Phos Adj Diet | Inotiv | TD.200790 |
|  | Iron deficient diet | Inotiv | TD. 09771 |
|  | Iron control diet | Inotiv | TD.09772 |
| In Vitro | BPGM siRNA | Horizon | L-008917-00-0005 |
|  | BPGM Human tagged ORF clone | OriGene | RG202105 |
|  | Recombinant Human FGF-23 Protein | R&D Systems | 2604-FG |
|  | Control siRNA | Horizon | D-001720-03-05 |
|  | *Calb1* cDNA | OriGene | MR203429 |
|  | *PCMV6* | OriGene | PS100001 |
|  | 2,3-Diphospho-D-glyceric acid pentasodium salt | Millipore Sigma | D5764-25MG |
|  | Deferoxamine mesylate salt | Millipore Sigma | D9533-1G |
|  | Seahorse XF Pyruvate | Agilent | 103578-100 |
|  | Seahorse XF Glucose | Agilent | 103577-100 |
|  | Seahorse XF DMEM | Agilent | 103575-100 |
|  | Seahorse XF DMEM Assay Pack | Agilent | 103680-100 |
|  | Seahorse XFe96/XF Pro *FluxPak* Mini | Agilent | 103793-100 |
|  | Seahorse XF Real-Time ATP Rate Assay Kit | Agilent | 103592-100 |
|  | 209/MDCT | ATCC | CRL-3250 |
|  | mKL-HEK293 | In-House Reagent | In-House Reagent |
|  | Eagle's Minimum Essential Medium | ATCC | 30-2003 |
|  | Fetal bovine serum | Corning | MT35016CV |
|  | Dulbecco’s Modified Eagle’s Medium | Gibco | 10-013-CV |
|  | Ham’s F-12 Nutrient Mix | Gibco | 11-765-054 |
|  | Penicillin-streptomycin | Millipore Sigma | P4333-100ML |
| Molecular | FGF23 Intact ELISA | QuidelOrtho | 60-6800 |
|  | FGF23 C-Term ELISA | QuidelOrtho | 60-6300 |
|  | PTH ELISA | QuidelOrtho | NC0195335 |
|  | KL antibody | Proteintech | 28100-1-AP |
|  | Actin antibody | Cell Signaling | 4970S |

## Notes

### Competing Interest Statement

The authors have declared no competing interest.

### Summary of Updates

NIH funding in the acknowledgment section.

